# Morphodynamics of human early brain organoid development

**DOI:** 10.1101/2023.08.21.553827

**Authors:** Akanksha Jain, Gilles Gut, Fátima Sanchis-Calleja, Ryoko Okamoto, Simon Streib, Zhisong He, Fides Zenk, Malgorzata Santel, Makiko Seimiya, René Holtackers, Sophie Martina Johanna Jansen, J. Gray Camp, Barbara Treutlein

**Author notes:** Correspondence: Akanksha Jain, J. Gray Camp and Barbara Treutlein. Contributed equally (Akanksha Jain, Gilles Gut).

## Abstract

Brain organoids enable the mechanistic study of human brain development, and provide opportunities to explore self-organization in unconstrained developmental systems. Here, we establish long-term, live light sheet microscopy on unguided brain organoids generated from fluorescently labeled human induced pluripotent stem cells, which enables tracking of tissue morphology, cell behaviors, and subcellular features over weeks of organoid development. We provide a novel dual-channel, multi-mosaic and multi-protein labeling strategy combined with a computational demultiplexing approach to enable simultaneous quantification of distinct subcellular features during organoid development. We track Actin, Tubulin, plasma membrane, nucleus, and nuclear envelope dynamics, and quantify cell morphometric and alignment changes during tissue state transitions including neuroepithelial induction, maturation, lumenization, and brain regionalization. Based on imaging and single-cell transcriptome modalities, we find that lumenal expansion and cell morphotype composition within the developing neuroepithelium are associated with modulation of gene expression programs involving extracellular matrix (ECM) pathway regulators and mechanosensing. We show that an extrinsically provided matrix enhances lumen expansion as well as telencephalon formation, and unguided organoids grown in the absence of an extrinsic matrix have altered morphologies with increased neural crest and caudalized tissue identity. Matrix-induced regional guidance and lumen morphogenesis are linked to the WNT and Hippo (YAP1) signaling pathways, including spatially restricted induction of the Wnt Ligand Secretion Mediator (WLS) that marks the earliest emergence of non-telencephalic brain regions. Altogether, our work provides a new inroad into studying human brain morphodynamics, and supports a view that matrix-linked mechanosensing dynamics play a central role during brain regionalization.

## Introduction

Unguided human neural or brain organoids generated from pluripotent stem cells (PSCs) develop self-organized regionalized domains composed of cell types and states with remarkable structural and molecular similarities to primary tissue counterparts^1−3^. Unguided brain organoid development proceeds through assembly, self-patterning, and morphogenetic mechanisms that reflect a latent intrinsic order emerging from the initial conditions of the system^4^. Multi-potent embryoid bodies (EBs) are directed towards neuroectoderm and the developing tissue can be supplied with an extrinsic matrix, such as matrigel, that supports formation and expansion of a polarized neuroepithelium surrounding large luminal regions^5−7^. Regional domains form harboring different neural progenitor cell states that develop, proliferate and ultimately differentiate into diverse neuronal cell types^8^. Extracellular matrix (ECM) proteins and glycoproteins such as Laminin, Decorin, and HAPLN1 are involved in many aspects of brain development^9^, and can be secreted from various cell types within and surrounding the developing brain (e.g. neural progenitor cells, meningeal cells) thereby modifying extracellular microenvironments^10,11^. Much of what is known about ECM secretion and its role in brain development derives from studies in non-human model systems, and it has remained unclear how the extracellular microenvironment impacts the early stages of human brain development. Organoid protocols exist that guide the development of specific brain regions by providing patterning molecules (morphogens such as BMP, SHH, FGF, SHH, etc.) to the culture media^12−14^ and some of these protocols do not use an extrinsic extracellular matrix for the initial neuroectoderm formation^15,16^. It has been difficult to understand early brain organoid morphodynamics and the role of extracellular microenvironment in shaping organoid morphogenetic patterning due to the lack of methods to dynamically track organoid development over many days. Recent advances in CRISPR-based stable fluorescent reporter tagging in stem cells and light sheet microscopy are providing opportunities for in toto imaging of fluorescently-labeled *in vitro* self-organizing systems^17−20^.

Current human brain organoid protocols pose challenges for live imaging because the organoids are relatively large in size, optically dense, slow in their development, and require sterile imaging conditions for weeks to months of development. Here, we address these challenges by developing a protocol for the generation of multi-mosaic, sparsely labeled brain organoids that are amenable to long-term live imaging, tracking, and segmentation. We use this protocol, together with long-term light sheet microscopy and a suite of computational tools, to study tissue morphodynamics, cellular behaviors, and interactions with ECM over two weeks of organoid development. We quantify cell morphologies as PSCs transition into a pseudostratified neuroepithelium, and observe interkinetic nuclear migrations, elongation of radial glial cells, and differentiation into neurons. We find that exposure to an extrinsic extracellular matrix (Matrigel) modulates tissue morphogenesis by inducing cell polarization and neuroepithelial formation, fostering lumen enlargement, and altering the global patterning and regionalization of the organoids. These changes in tissue patterning are associated with modulation of the WNT signaling pathway, and in particular YAP mediated upregulation of Wnt Ligand Secretion Mediator (WLS) expression. Altogether, we establish a multiscale morphodynamic view of human brain organoid formation.

### Long-term live imaging of sparse and multi-mosaic fluorescently labeled brain organoids

We established a protocol to generate sparse, mosaically-labeled, fluorescent, brain organoids that are amenable to long-term imaging using light sheet fluorescence microscopy (Fig. 1a). Induced PSCs (approximately 500 cells, see methods and Supplementary methods table 1) containing genetic fluorescent labels are aggregated at day 0 into spherical embryoid bodies (EBs) and cultured in media maintaining proliferation and multipotency until day 4, when the organoids are transitioned into neural induction media (NIM) containing extrinsic matrix (matrigel). At day 10, media is exchanged to enhance neural differentiation, and at day 15, vitamin A is provided to support maturation. Compared to a previously published unguided brain organoid protocol^1,21^, the lower number of input cells and early exposure to matrix and neural induction leads to organoids with a smaller initial size and earlier expansion of lumina surrounded by neuroepithelium (Extended Data Fig. 1a-c). Time course single-cell transcriptomics (day 5, 7, 11, 16, 21) revealed transitions from neuroectodermal progenitor states (days 5-11) via early prosencephalic neural progenitor states to regionalized neural progenitor states (days 11-21) of predominantly telencephalon and diencephalon identity (Fig. 1b-e). Whole-mount fluorescent *in situ* hybridization chain reaction (HCR) revealed spatial segregation of these developing brain regions (Fig. 1f; Extended Data Fig. 2).

**Fig. 1.**
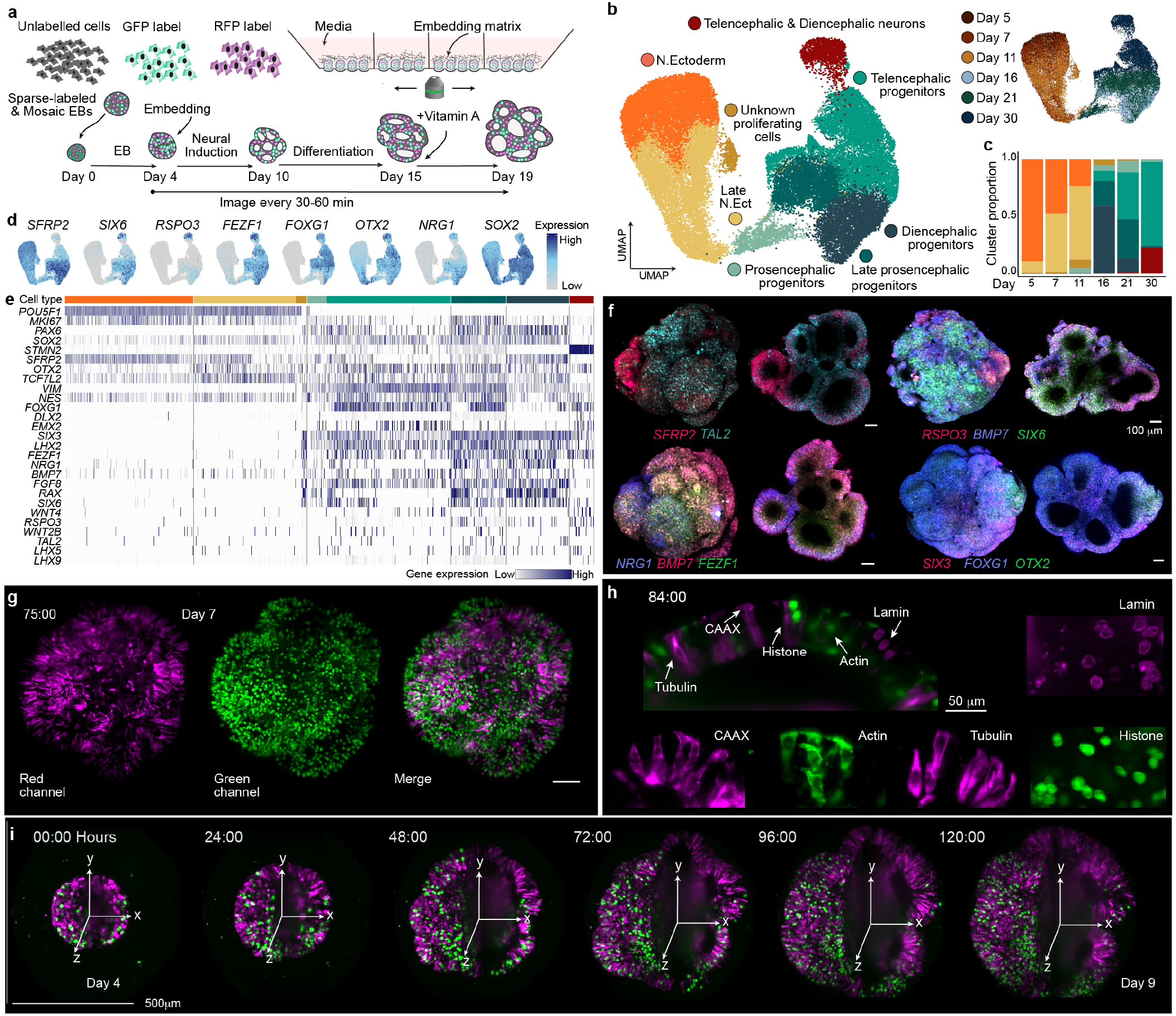
Long-term live imaging of sparse and multi-mosaic fluorescently labeled brain organoids. a) Schematic of the mosaic fluorescent organoid protocol and light sheet image acquisition setup. b) UMAP embedding of organoid time-course scRNA-seq data with cells colored by cluster and labeled by cell population (left) or time point (right). c) Stacked barplot showing the proportion of each cell population per time point. d) Feature plot showing normalized expression of representative marker genes. e) Heatmap showing normalized marker gene expression grouped per cluster. f) 3D projections (left) and cross sections (right) of 4 different organoids (Day 15) stained for marker transcripts with whole-mount fluorescence *in situ* hybridization chain reaction (HCR). Scale, 100 micrometer. g) Maximum projection image at 75 hours of imaging from a 188 hour imaging experiment. Organoids contain 5 different cell lines that contain stable genetic tagging of proteins with red or green fluorescent protein (RFP, GFP), as well as unlabeled cells. Scale, 100 micrometer. h) Organoid cross section (84 hour) showing nuclear membrane (Lamin, RFP, magenta), plasma membrane label (CAAX, RFP, magenta), Actin (GFP, green), Tubulin (RFP, magenta), and nuclei (Histone, GFP, green). i) Images at different time points showing the maximum intensity projection (left half) and cross section (right half).

For live imaging experiments, day 4 organoids are moved to the imaging chamber, covered with matrix to stabilize tissue location, and provided with the neural induction medium (NIM) (Fig. 1a; Extended Data Fig. 1d). This protocol enables light sheet imaging for weeks of development using an inverted light sheet platform with controlled environmental conditions suitable for *in vitro* cell culture applications (Viventis Microscopy sárl). The custom microscope is adapted for long-term brain organoid imaging with a 25x objective demagnified to 18.5x with a 710*µ*m field of view that captures the entire organoid during the first week of development, followed by tiling acquisition as the organoids grow larger (Extended Data Fig. 1e). We adapted the sample mounting chamber to allow for stable long-term imaging over weeks, enabling media exchanges with limited drift (Extended Data Fig. 1f,g). The custom sample chamber is composed of fluorinated ethylene propylene (FEP) bottom with rounded cone pockets of 800*µ*m diameter such that one organoid is added per microwell. The sample chamber is divided into 4 sub-chambers with vertical walls to separate different imaging conditions, in total containing 4 microwells per subchamber and enabling parallel imaging of up to 16 organoids in one experiment for 1-3 weeks (Extended Data Fig. 1d).

To explore cellular dynamics during organoid development, we used a set of iPSC lines (based on WTC-11)^22^ each expressing a single endogenously tagged protein representing a particular organelle or cellular structure including the plasma membrane (CAAX, RFP), Actin cytoskeleton (Actin (ACTB), GFP), microtubules (Tubulin (TUBA1B), RFP), nucleus (Histone (HIST1H2BJ), GFP), and nuclear envelope (Lamin (LAMB1), RFP). We combined these 5 labeled lines together with the unlabeled parental WTC-11 line at a ratio of 2:100 (labeled:unlabeled) to achieve sparse mosaicism for tracking and resolving single nuclei/cells for segmentation. This enables profiling of the dynamics of multiple subcellular features in a single cross sectional area of the developing organoids (Fig. 1g,h). Starting at day 4, organoids were imaged for 188 hours with a 30 minute time resolution to track one week of organoid development. This imaging time frame follows organoids as they transition from spherical EBs to form a neuroepithelium composed of expanding lumina that begin self-patterning at around 2 weeks (Fig. 1i; Supplementary video 1-2). Altogether, this approach provides a framework for imaging and exploring multiple phases of organoid development spanning neuroepithelial formation and brain regionalization.

### Extrinsic extracellular matrix impacts brain organoid morphogenesis

To quantify morphodynamic variation across organoids, we imaged 16 organoids in parallel in a single imaging experiment (Fig. 2a; Extended Data Fig. 3). After approximately 24 hours of imaging, on day 5, we observed multiple cavitation spots in each organoid, which over time expanded into lumina surrounded by neuroepithelium (Fig. 2b; Supplementary video 3). We segmented and quantified tissue scale properties such as organoid volume, lumen volume, and lumen number per organoid to assess the tissue morphodynamics associated with neural induction, neuroepithelium formation and patterning of the organoids (Fig. 2c; Extended Data Fig. 3a,b; Supplementary video 4-5). Despite qualitative differences in tissue morphologies and organoid growth, we found that all 16 organoids exhibited consistent growth dynamics. Between day 4 and 8, organoids experienced a 4-fold increase in overall volume (Fig. 2d), accompanied by an increase in total lumen volume from day 5 to day 8 (Fig. 2e; Extended Data Fig. 3c). The average lumen number per organoid first increased from 3.7 ± 2.5 to 13.4 ± 2.5 between day 5-6 and then decreased again to an average number of 5.4 lumina per organoid, indicating fusion of small lumen (Fig. 2f,g). After day 7, the lumen number per organoid remained stable whereas the lumen volume decreased corresponding to an increase in neuroepithelial thickness (Fig. 2f,g; Supplementary video 6). These observations highlight three morphodynamic phases of early brain organoid development including an early phase of rapid tissue and lumen growth, a phase of tissue stabilization involving lumen fusion events and finally a phase of neuroepithelium maturation. To understand the cell state changes associated with these tissue transitions, we subsetted the scRNAseq data from day 5, 7 and 11 organoids, established a diffusion component-based pseudotemporal ordering of cells and explored pseudotime-dependent expression changes (Fig. 2h-i). We identified a major transcriptomic switch resolving the cell state transition from an early neuroectoderm-like progenitor state (POU5F1, ITGA5, PROM1, THY1) to a later neural tube-like neuroepithelial progenitor state (SOX2, TCF7L2, OTX2, ZIC2, CYP26A1, ITGA6, SOX21) (Fig. 2j; Supplementary Table 1,2). Genes that were upregulated over pseudotime showed a Gene Ontology (GO) enrichment for Non-motile cilium, Myosin complex, and ECM-related terms (e.g. Basement membrane, Collagen Trimer, Extracellular matrix) (Fig. 2k-l; Supplementary Table 2). Interestingly, genes upregulated over pseudotime included several ECM proteins (COL1A1, COL11A1, LAMA5) and ECM interactors (ITGA6, HAPLN3, MMP16, IGFBP2), some of which have been detected in primary neural progenitor cells^11,23,24^. To understand the impact of an extrinsically provided basement membrane rich matrix on the observed tissue and cell state transitions, we compared development of organoids cultured with matrigel as an extrinsic matrix with organoids cultured without any extrinsic matrix or embedded in 0.6% low melting agarose (Fig. 2m; Extended Data Fig. 6a,b). Agarose is a polysaccharide gel that should provide an inert diffusion barrier to capture some of the secreted ECM and other patterning molecules (morphogens) intrinsically produced from the organoid neuroepithelium^25^. Organoids grown without matrigel were smaller and had different tissue morphologies lacking outgrowing neuroepithelium buds (Fig. 2m; Extended Data Fig. 3d). We used our long-term light sheet microscopy setup to follow the organoid morpho-dynamics and found that in all conditions organoids developed lumina, but the overall lumen volume and lumen growth dynamics differed between the conditions (Fig. 2n-p; Extended Data Fig. 3d; Supplementary video 7). Organoids grown without any extrinsic matrix showed a slower growth of overall organoid and lumen volume with a maximum over-all lumen volume at day 8 in contrast to day 6.5 for organoids grown in the presence of matrigel (Fig. 2q-s; Extended Data Fig. 3e). Interestingly, in the presence of an inert external diffusion barrier (agarose), lumen expansion was also delayed compared to the matrigel counterpart, but less than for the organoids without matrix. Immunohistochemistry confirmed that the basal ECM component COL4A1 accumulates around the periphery of the organoid, and the apical marker CDH2 showed a restricted expression around the inner cavity walls highlighting that lumina in the organoids exhibit apicobasal polarity induced by matrigel. However in organoids grown without any extrinsic matrix neuroepithelium polarity is either inverted with secretion of COL4A1 inside the lumen cavities or is mixed with both inner or peripheral accumulation of CDH2 in outer or inner walls of different lumen. (Fig. 2t). Altogether, these data showed that the presence of extrinsic extracellular matrix has a major impact on tissue-scale morphogenesis in human brain organoids and indicated that altered tissue polarities might impact cellular morphodynamics in the maturing neuroepithelium.

**Fig. 2.**
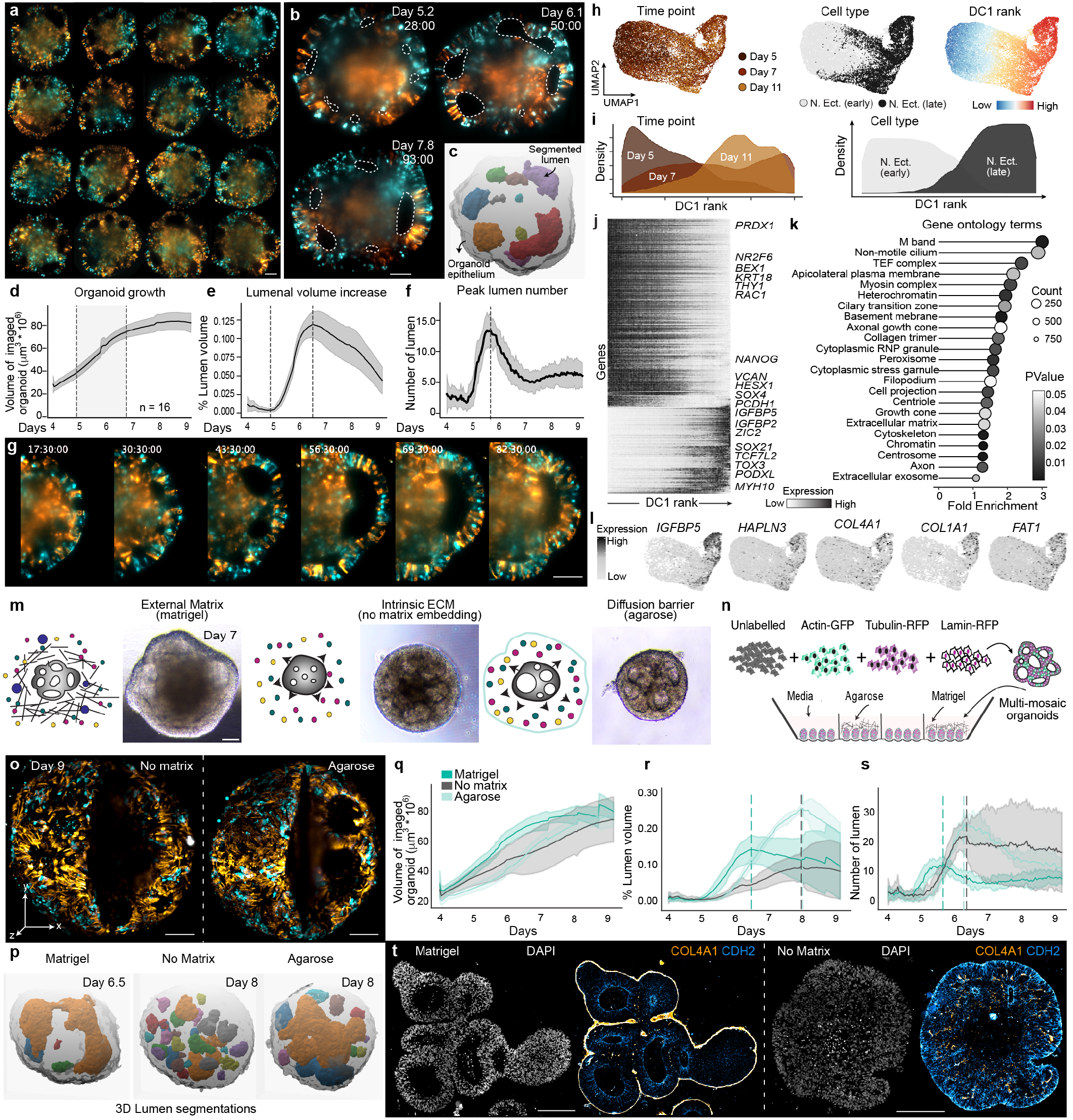
Exogenous extracellular matrix impacts brain organoid morphogenesis. a) Organoid projections (Day 7) for 16 simultaneous image acquisitions showing nuclear membrane (Lamin, RFP, orange), plasma membrane label (CAAX, RFP, orange), Actin (GFP, cyan), Tubulin (RFP, orange), and nuclei (Histone, GFP, cyan). Maximum projection (left) and cross section (right). Scale, 100 micrometer. b) Stills at different time points from the same organoid showing cross section and highlighting lumenal morphologies (dashed lines). c) Cross section of an organoid showing segmented lumen and organoid epithelium masks. d) Graph showing total organoid volume measured per day from Day 4-9. Error bars indicate standard deviation (d-f). e) Graph showing total volume of all lumen over time. f) Graph showing change in total number of segmented lumen over time. g) Cross sections of an organoid showing lumen formation and fusion over time. h) UMAP embedding of scRNA-seq data with cells colored by day, cell type and diffusion component ranking. i) Plot showing distribution of cells arranged in a pseudotime prediction and showing the composition changes of neuroectoderm and neural epithelium. j) Heatmap showing normalized gene expression over diffusion component ranking. k) Top DAVID GO analysis terms calculated for genes that change over pseudotime from day 5 to day 11. l) Feature plot showing normalized expression of example ECM-related genes that show an increase in expression over time. m) Schematic representation of the extracellular microenvironment and the corresponding brightfield image for organids grown with Matrigel (External ECM), without any external embedding (Internal ECM) and with a low melting agarose embedding (Diffusion barrier). n) Schematic showing the light sheet experimental setup to acquire simultaneous imaging of 16x organoids with different embedding treatments. o) Image stills at Day 9 showing the maximum projection (left) and cross section (right) of an organoid grown without any embedding and with an agarose diffusion barrier. p) 3D renderings of segmented lumen in organoids grown with matrigel, with no matrix and with a diffusion barrier. q-r) Graphs showing total organoid volume measured per day from day 4-9 for all imaged organoids (q), change in the total number of segmented lumen (r), and total volume of all lumen over time (s). Scale, 100 micrometers. Organoids imaged per condition, 4. Error bars indicate standard deviation. t) Images show cross sections from an immunohistochemistry experiment on organoids (day 15) grown with matrigel or with no matrix and stained to show nuclei (DAPI), COL4A1 and CDH2. Scale Bar is 100 micrometers in all images.

### Single-cell morphotype analysis reveals shape transitions within developing organoids

We leveraged our multi-mosaic labeling of plasma membrane, Actin, Tubulin, nuclear membrane and Histone to assess the impact of extracellular matrix on cell morphology changes during early brain organoid development. We developed an image analysis pipeline that uses spatial embedding-based instance segmentation^26^ to predict cell and nucleus masks, and used morphometric feature extraction and a random forest classifier to demultiplex the fluorescence signals into the 5 individual labeled structures (Fig. 3a-c; Extended Data Fig. 4). We generated a UMAP^27^ embedding based on extracted morphological features with each point representing one cell structure. This revealed two major groups of nuclear (Histone, Lamin) and cellular (Actin, Tubulin, membrane) structures, respectively (Fig. 3d). To quantify structural changes in the developing organoid, we performed high-resolution clustering to group structures into morphotypes (we define a morphotype as a cluster of cells with similar morphological features such as volume, curvature, axis length), and used PAGA^28^ trajectory analysis to identify a gradient of structural changes in the tissue (Fig. 3e). This revealed a time-dependent cytoskeletal and membrane elongation as well as compression of nuclei (Fig. 3e; Extended Data Fig. 4). Detection of the major cell axis and quantification of a cell alignment index (cosine angle to the organoid surface) showed that cells align perpendicular to the organoid surface after exposure to matrigel (Fig. 3f; Extended Data Fig. 4). Together, these analyses quantify the transition of multipotent stem cells to form an early neuroectoderm, which changes into a matured neuroepithelium based on cell structure morphology and tissue topology.

**Fig. 3.**
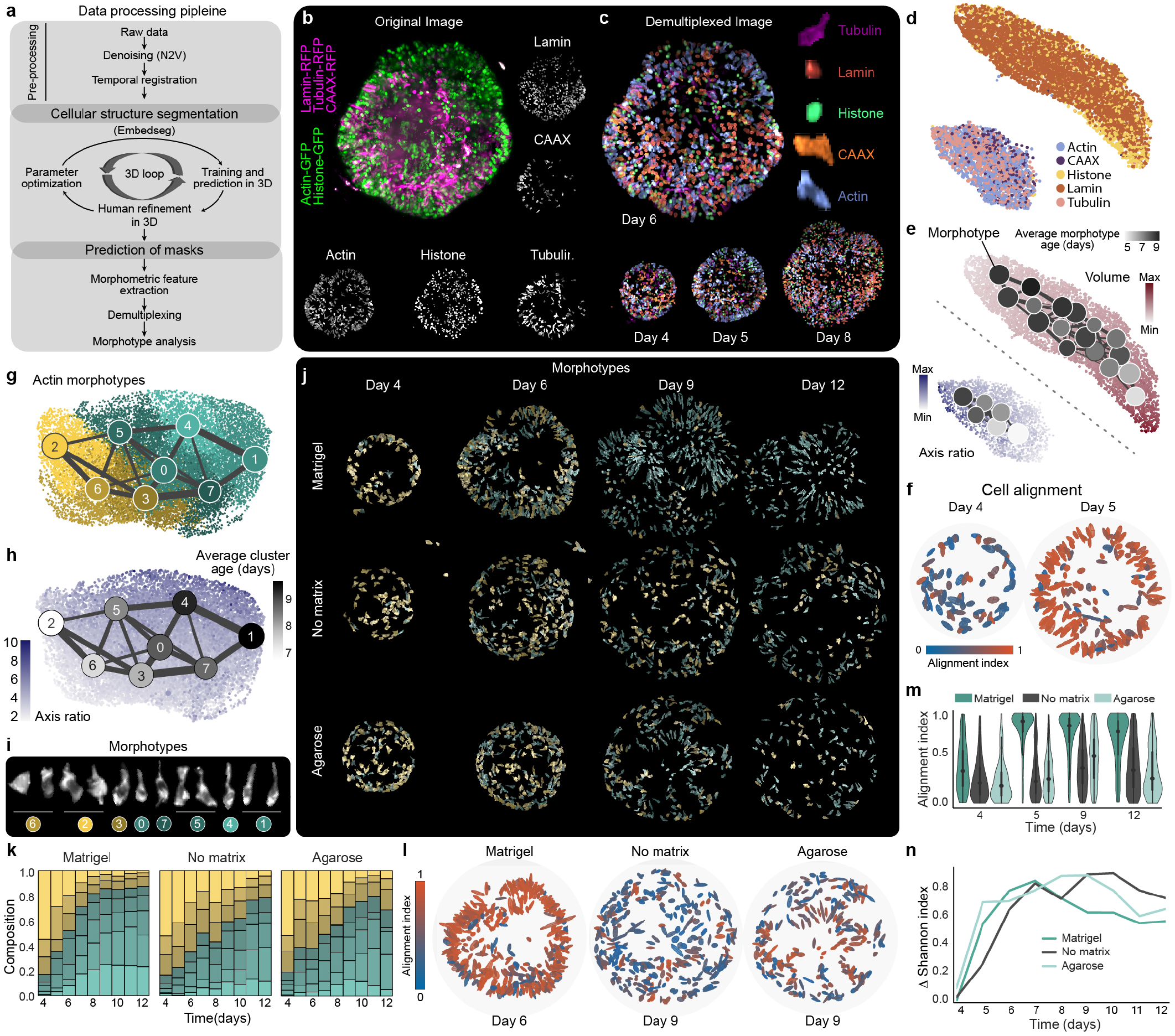
Cell and nuclei morphology transitions using demultiplexed mosaic cellular labels. a) Schematic overview of the analysis strategy used for cell segmentation and demultiplexing of mosaic cell labels. b) Maximum intensity projection of an organoid at day 6 labeled with nuclear membrane (Lamin, RFP, pink), plasma membrane (CAAX, RFP, pink), Actin (GFP, green), Tubulin (RFP, pink), and nuclei (Histone, GFP, green). Scale, 100 micrometer. c) Demultiplexed image (Lamin, dark brown; CAAX, orange; Actin, blue; Tubulin, magenta; Histone, green). Scale, 100 micrometer. d) PAGA initialized UMAP embeddings of all demultiplexed labels based on morphometric feature extraction. e) PAGA initialized UMAP embeddings showing change in axis length for Actin, Tubulin and CAAX labels and change in nuclei volume measured using Histone and Lamin segmentations. PAGA plots show change in average cluster age (days), node size indicates the number of cells within one cluster, and edge width reflects the strength of connection between two clusters. f) All cells (Actin) colored by their alignment index (absolute cosine angle to the nearest organoid surface normal) in matrigel condition. Scale is from 0-1 with 1 (red) corresponding to cells that align perpendicular to the organoid surface. g) PAGA initialized UMAP embeddings and PAGA plots showing cell morphotype clusters using cells segmented from matrigel, no matrix, and agarose conditions. The plots are based on morphometric measurements extracted for all segmented cells (Actin). h) PAGA initialized UMAP embeddings shows change in axis ratio of cells over time overlaid with average cluster age shown using PAGA plots. PAGA plots are color-coded based on the average age of the cluster from light gray to black. i) Example cells (Actin) belonging to each of the morphotype clusters. j) Example images of organoids with Actin labeled cells showing cells colored by their morphotype clusters. k) Stacked bar plots showing proportion of cells in individual Actin morphotype clusters in matrigel, no matrix and agarose conditions. l) Image shows cells (Actin) colored for the cell alignment index in matrigel condition at day 6 and no matrix and agarose at day 9. m) Violin plot showing the cell alignment (Actin) values across all segmented cells from day 4 to day 12 for all three conditions. n) Line plot showing the change in Shannon index between the three conditions based on change in the number of cells per cluster over time (matrigel (n=3), no matrix (n=3) and agarose (n=3) in k.

We next applied this single-cell morphotype approach to assess the effect of matrix perturbation on cell shape transitions. We segmented labeled structures at each day of imaging from organoids in each matrix condition (matrigel, agarose, and no extrinsic matrix) and assessed morphotype heterogeneity focusing on cytoskeletal labels (Actin, Tubulin) (Fig. 3g; Extended Data Fig. 5a,b). We found 8 Actin morphotypes that aligned along a temporal gradient corresponding to increasing cell elongation over time (5 elongated morphotypes in cyan and 3 non-elongated morphotypes in yellow) (Fig. 3g-j). In the presence of matrigel, organoids exhibited elongated morphotype clusters at an earlier time point compared to organoids grown in the absence of matrigel (Fig. 3k). In addition, only a small proportion of cells were aligned perpendicular to the surface in organoids without matrigel (Fig. 3l,m; Extended Data Fig. 5c,d). Consistent with these observations, organoids grown without matrigel showed a higher Shannon index (a diversity index) reflecting a more heterogeneous population of morphotypes per day (Fig. 3n). We note that the presence of an inert diffusion barrier (agarose) induced higher cell alignment and cell elongation, and lower Shannon index than was seen in absence of matrigel (Fig. 3j-n). Together with the previous analysis of lumenal architectures, these data show that organoids generated without matrigel as an external ECM have alterations in tissue topology, contain a larger proportion of non-aligned and non-elongated cells, show higher heterogeneity in cell morphotypes and do not form a homogeneous neuroectoderm and neuroepithelium.

### Matrix impacts organoid morphogenesis and patterning through WNT pathway modulation

To understand the molecular cell states developing in organoids grown under the different matrix conditions (matrigel, agarose, no matrix), we performed single-cell transcriptome analysis of day 13 organoids (Fig. 4a,b; Extended Data Fig. 6a-f). We identified 8 cell populations including telencephalic and non-telencephalic neural progenitors, as well as neural crest and peripheral nervous system neurons. The proportions of each population differed between matrix conditions, with organoids grown in the presence of matrigel containing significantly more telencephalic progenitors and less neural crest cells than no-matrix or agarose counterparts (Fig. 4a,b). The gene ontology analysis on differentially expressed genes between matrigel and no matrix conditions showed enrichments for multiple signaling pathways, including components of WNT, Notch, FGF, and Hippo signaling pathways, as well as genes associated with Actin cytoskeleton regulation (Fig. 4c; Supplementary Table 3-6). Consistent with the enriched proportion of telencephalic progenitors, multiple transcription factors associated with anterior fates of the developing neural tube (e.g. SIX3, LHX2, NRG1, FOXH1, HESX1) were expressed higher in matrigel-exposed organoids. Interestingly, we identified WLS, RSPO3 and GPC3, as highly upregulated in organoids grown without matrix, indicating that the Wnt/beta-catenin signaling pathway might be upregulated in this condition (Fig. 4c). WLS has been previously identified as one of the earliest markers of non-telencephalic fate in human brain organoids^3^, and WLS as well as other WNT-associated genes upregulated in the nomatrix condition are highly expressed in non-telencephalic cell populations in the primary human developing brain^29^ (Fig. 4c).

**Fig. 4.**
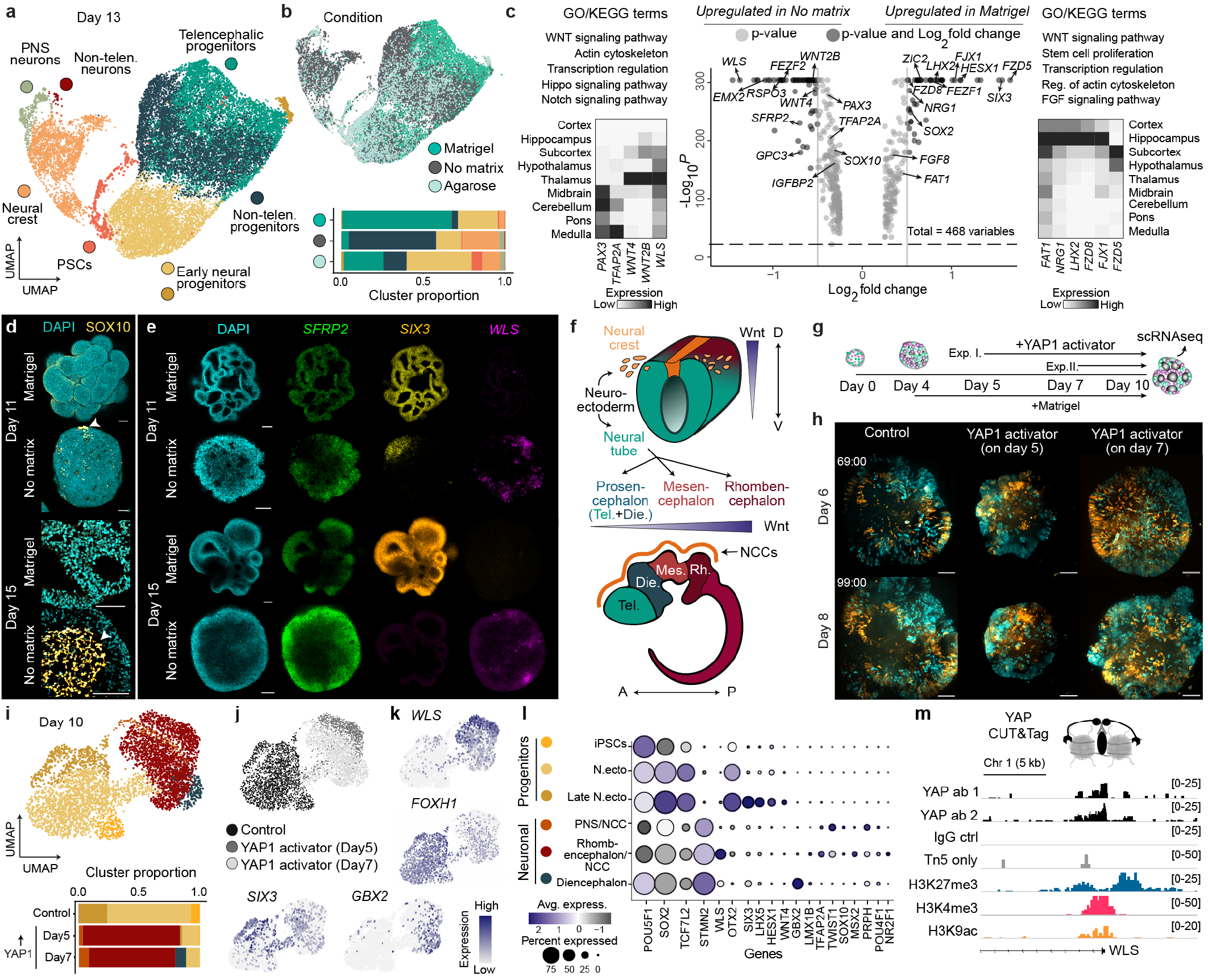
Matrix impacts organoid morphogenesis and patterning through WNT pathway modulation and YAP mechanotransduction. a) UMAP embedding of scRNA-seq data with cells colored by cell population and stacked barplot showing the cluster proportion of each cell population per treatment. b) UMAP embedding of scRNA-seq data with cells colored by treatment. c) Volcano plot showing differentially expressed genes upregulated in matrigel (right) and upregulated in No matrix (left). GO/KEGG terms for genes differentially upregulated between matrigel and No matrix conditions. Heatmaps show average gene expression (of selected differentially expressed genes between matrigel and no matrigel) in different regions of the human fetal brain tissue from 5-14 post-conceptional weeks (data sourced from^29^). d) Maximum intensity projection images from an immunofluorescence staining of whole mount fixed organoids (day 11) showing neural crest cells marked by SOX10 antibody (arrowhead) and nuclei labeled with DAPI and cross sections at day 15. e) Cross sections from a whole mount fluorescence *in situ* hybridization chain reaction (HCR) staining for selected differentially expressed gene between matrigel and no matrix on organoids day 11 and day 15, Scale, 100 micrometer. f) Schematics showing the tissue state transitions in brain development and the putative expression of Wnt signaling genes in the developing neural tube. Top: neuroectoderm forms the neural tube and the neural crest cells. Bottom: neural tube forms the primary brain vesicles, prosencephalon (telencephalon (Tel.) + diencephalon (Die.)), mesencephalon (Mes.) and rhombencephalon (Rh.). Orange marks the neural crest cells (NCCs). g) Schematic overview of the light sheet imaging and scRNAseq experiment with control and YAP1 activator treated organoids. All EBs were given neural induction media with matrigel embedding on day 4. Experiment I. YAP1 activator (Py-60) was added to the imaging subchamber on day 5, Experiment II. YAP1 activator (Py-60) was added to the imaging subchamber on day 7. Control organoids were given DMSO. Imaging was stopped on day 10 and corresponding organoids from all three conditions grown in the cell culture incubator were profiled with scRNAseq on day 10. h) Maximum intensity projections on day 6 and day 8 of representative organoids. Sparse and multi-mosaic organoids contain cells labeled with nuclear membrane (Lamin, RFP, orange), Actin (GFP, cyan) and Tubulin (RFP, orange) and unlabelled cells. Scale, 100 micrometer. i) UMAP embedding of scRNA-seq data from day 10 organoids in control and YAP1 treatment conditions. Colors show regional cell populations and stacked barplot showing the cluster proportion of each cell population. j) UMAP embedding of scRNA-seq data with cells colored by treatment (control, YAP1 activator given on day 5, YAP1 activator given on day 7). k) Feature plots showing expression of selected genes in the UMAP embedding. l) Dotplot showing average expression and percentage of cells expressing selected regional marker genes per cell populations in i. Abbreviations: N.ecto = neuroectoderm, NCC = neural crest cells, PNS = peripheral nervous system. m) Genome browser snapshots of a bulk CUT&Tag experiment showing the enrichment score of YAP1 binding to the WLS gene, profiled with two different YAP1 antibodies, IgG and Tn5 control, and the repressive and active marks profiled with H3K27me3, H3K4me3 and H3k9ac antibodies.

Immunostaining confirmed a higher proportion of neural crest cells (SOX10) emerge in organoids without matrix (Fig. 4d), which supports the presence of a non-polarized morphotype observed in the light sheet imaging data. Whole-mount HCR staining for WLS, SIX3, and SFRP2 confirmed the differential expression of these genes between matrigel and no-matrix conditions (Fig. 4e; Extended Data Fig. 6g, Extended Data Fig 7). Wnt signaling is known to regulate rostral-caudal patterning of the neural tube, and neural crest differentiation at the neural plate border^30−32^. Together, these analyses indicated that the dorso-ventral and rostral-caudal axes are altered by the addition of matrigel (Fig. 4f).

Due to the strong morphological changes in the neuroectoderm induced by matrigel, as well as the upregulation of the WNT and Hippo signaling pathways in no-matrix conditions, we tested the effect of YAP modulation on organoid development. YAP1 has been reported to upregulate WNT pathway genes such as WLS in cardiomyocytes^33,34^ and mediate crosstalk between WNT and Hippo pathways^35^. We activated the YAP1 pathway by providing Py-60^36^ to the organoid culture at different, early time points (day 5, 7 or 10) and analyzed organoids using scRNAseq at day 10 (5 and 3 days of activation, respectively) and day 16 (6 days of activation) of organoid development, respectively (Fig. 4g; Extended Data Fig. 6; Supplementary video 10). Organoids that were grown in matrigel with YAP1 over-activation exhibited an inability to expand lumina (when provided on day 5) or showed a collapse of lumina (when provided on day 7) resulting in altered organoid and lumen morphologies (Fig. 4h; Extended Data Fig. 6h). Single-cell transcriptome analysis revealed a strong effect of YAP1 activation with cells from control and treatment conditions largely segregating in the UMAP embedding and composing different clusters (Fig. 4i-l; Extended Data Fig. 6i-l). Genes upregulated upon YAP1 activation included WLS, TFAP2A, MSX2 and PRTG, whereas genes expressed higher in control cells included ZIC2, OTX2, IGFBP2 and FOXH1 (Fig. 4k,l; Supplementary Table 7,8). In addition, we found expression of GBX2, LMX1B, PRPH and STMN2 in YAP1 activation condition, consistent with caudalization of brain organoid tissue and early neurogenesis (Fig. 4l; Extended Data Fig. 6l). Finally, through a bulk CUT&Tag experiment in the neuroepithelium, we show that YAP1 directly binds the WLS promoter region (Fig. 4m). Taken altogether, we show that application of matrigel, an external basement membrane-rich ECM, leads to lumen expansion and self-patterning into largely rostal prosencephalic regions, and that activation of YAP1 and increased WLS expression levels promote caudalization of the developing neuroepithelium.

## Discussion

Understanding the developmental dynamics of mammalian neural tube morphogenesis and patterning, corresponding to the first few days of mouse embryogenesis and 2-5 weeks of human embryonic development, have remained obscure due to challenges in accessing live neural tissue. Brain organoids enable modeling important aspects of human neuroepithelium morphogenesis *in vitro*, however, there have been technical challenges that have inhibited insight into the dynamics of how neural tissues self-organize, including the opaque nature of the tissue and long developmental times. Here, we overcome these obstacles, harnessing the modularity inherent to brain organoid protocols to generate sparsely labeled, multi-mosaic organoids. This strategy enables incorporation of multiple fluorescent signals that can be imaged at high spatiotemporal resolution for several days leveraging the low phototoxicity offered by light sheet microscopy^37,38^. We achieve sterile growth conditions that enable long-term tracking of cellular and subcellular signals at single-cell resolution, and developed a computational pipeline to identify structural morphotypes using this imaging modality. Our analysis pipeline streamlines preprocessing and post-processing of raw image data to generate segmented lumen and organoid masks to track tissue level morphological changes, and to generate cell masks to allow quantification of cell shape morphometrics and alignments from hundreds to thousands of cells spanning an entire week of organoid development. Using this system, we have generated a detailed characterization of the cell and tissue dynamics in unguided brain organoid development from induction of the neuroectoderm to formation of a patterned neuroepithelium that gives rise to distinct brain regional progenitors. The extracellular matrix and the mechanical environment have been postulated to play a role in both the morphogenesis and patterning of the neural tube; however, the role of an externally provided matrix in neuroepithelium organization has remained unclear.

Using our long-term live imaging framework in combination with single-cell transcriptomic analyses, we demonstrate that extracellular matrix provided on the outer margin of organoids induces stem cells to efficiently form the neuroectoderm with polarized cells that arrange perpendicular to expanding, ventricle-like lumina and eventually influences patterning of the tissue into largely telencephalic domains. In the absence of an extrinsic matrix, organoids show mixed cell alignments without apico-basal polarity oriented in an inner-outer axis around the lumen, resulting in accumulation of the secreted ECM in both the lumen cavity and outside of the organoid. We find that canonical WNT and YAP1/Hippo pathways are involved in this matrix-mediated neuroepithelium alteration, which has an influence on both rostral-caudal and dorso-ventral brain organoid patterning. Ectopic YAP1^39^ activation also led to organoid caudalization with an increase in non-telencephalic cell types, which was associated with loss of lumen expansion. Interestingly, WLS and other WNT pathway-related genes were similarly modulated under no-matrix and YAP1 activation conditions, suggesting that YAP1 plays a role during both matrix-mediated morphogenesis and organoid patterning. Together, our work provides a technological advance towards understanding the morphodynamics of organoid development, provides mechanistic insights into matrix-mediated neuroepithelial signaling pathways, and paves the way for future explorations of the extracellular microenvironment during human brain development.

## Supporting information

Supplementary material guide

Supplementary videos guide

## AUTHOR CONTRIBUTIONS

A.J. generated the organoids used in this study with support from R.O., F.S.C., S.S and S.M.J.J. A.J. acquired the light sheet, HCR and immunohistochemistry imaging data used in this study. A.J generated the single-cell transcriptome datasets with support from M.Santel, F.S.C and M.Seimiya. A.J performed the HCR and immunohistochemistry experiments with support from S.S and M.Seimiya. A.J performed the single-cell transcriptome data analysis with support from G.G, image analysis of HCR and immunohistochemistry datasets, and light sheet data analysis. Z.H performed the pseudotime analysis and primary human brain data analysis. A.J and R.H performed the iterative immunohistochemistry. F.Z. performed the bulk CUT&Tag experiment. G.G performed the light sheet data analysis with organoid tissue segmentation and quantification analysis, developed the cell demultiplexing analysis pipeline, organoid alignment analysis, and morphotype analysis with support from A.J. A.J, B.T. and J.G.C. designed the study and A.J., G.G., B.T. and J.G.C. wrote the manuscript.

## DATA AVAILABILITY

Raw sequencing data will be deposited into European Genome Phenome Archive (https://ega-archive.org/). Processed scRNAseq data will be made available via Zenodo. Due to its large size the light sheet data will be made available upon request. All experimental materials are available upon request to.

## CODE AVAILABILITY

All code generated in the study including analysis parameters will be available at GitHub 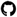(https://github.com)(quadbio/morphodynamics_human_brain_organoid) or upon request.

## ACKNOWLEDGEMENTS

We thank the members of the Treutlein and Camp laboratories, P. Liberali and G. de Medeiros for discussions; Petr Strnad, Andrea Boni and Viventis Microscopy for helping with set up of light sheet microscopy, Manan Lalit for support with EmbedSeg, Kevin Yamauchi and Christoph Harmel for discussions about morphometrics and imaging data analysis, P. Dubey, the Coriell Institute and the Allen Cell collection for the iPSC lines, staff at the Murdoch Children’s Research Institute and Murdoch University for providing the HES-3 NKX2-1:GFP cell line. Illumina sequencing was performed by I. Nissen, E. Vogel Burcklen and C. Beisel at the Genomics Facility at D-BSSE, ETH Zurich. Microscopy support was provided by E. Montani, J. C. Arias and T. Horn at the single-cell facility at D-BSSE, ETH Zurich. This work was supported by Chan Zuckerberg Initiative DAF, an advised fund of the Silicon Valley Community Foundation CZF2021-237566 (to J.G.C. and B.T.), the European Research Council (803441-Anthropoid, to J.G.C.; 758877-Organomics, to B.T.; 874606-Braintime, to B.T.), the Swiss National Science Foundation (project grant 310030_84795, to J.G.C.; project grant 310030_192604, to B.T.), the Swiss National Center of Competence in Research Molecular Systems Engineering (to B.T.).

## Materials and Methods

### Experimental methods

#### Stem cell and organoid culture

We used the following iPSC cell line for all experiments (also see Supplementary methods table 1) : Histone2B-mEGFP that uniformly labels nuclei (cell line ID: AICS-0061-036), mEGFP-Beta-Actin that uniformly labels ACTB (cell Line ID: AICS-0016 cl.184), mTagRFP–T–CAAX which labels cell membrane (cell line ID: AICS-0054-091), mTagRFP–T-Tubulin-alpha1b that labels TUBA1B (cell line ID: AICS-0031-035), mTagRFP–T-LaminB1 that labels LMNB1 (cell line ID: AICS-0034-062) and unlabeled WTC iPSCs (cell line ID GM25256). Stem cell lines were cultured in mTeSR™ Plus (Stem Cell Technologies) with mTeSR+ supplement (Stem Cell Technologies) and supplemented with penicillin–streptomycin (Pen-strep, 1:200, Gibco, 15140122) on Matrigel-coated plates (Corning, 354277). Cells were passaged 1–2 times per week using TryplE (Gibco, 12605010) or EDTA in DPBS (final concentration 0.5 mM) (Gibco, 12605010). The cell culture medium was supplemented with 1:200 Rho-associated protein kinase inhibitor (ROCKi) Y-27632 (final concentration 5 *µ*M, STEMCELL Technologies, 72302) on the first day after passage. All cells were tested for mycoplasma infection regularly using PCR validation (Venor GeM Classic, Minerva Biolabs) and found to be negative. The organoid generation protocol (multi-mosaic and sparse organoids) is as follows: 500 cells (except, Extended Data Fig. 1b with 3000 cells) in mTSR+ (with 1:200 ROCKi and 1:200 Pen-strep) were added per well of a 96 well plate (Corning, CLS7007) and centrifuged at 200g for 5 min to generate embryoid bodies (EBs). Fresh mTSR+ with 1:200 Rocki and 1:200 Pen-strep was exchanged on day 2. Fresh Neural induction media with 2% dissolved matrigel was supplied on day 4 and exchanged every other day, followed by differentiation media -VitA with 2% matrigel on Day 10 and differentiation media + VitA and with 1% matrigel on Day 15. Organoids were moved to 24-well plates on day 15, one organoid per well, and moved to a shaker, followed by moving one organoid per well to a 6-well plate at one month and kept on a shaker. The no matrix organoids were cultured following the exact same conditions, without any addition of matrigel at any point. For agarose embedding, organoids were embedded in 0.6% low melting agarose (SeaPlaqueTM Agarose, 501010), stock is 1% in PBS, diluted to 0.6% in neural induction media. The composition of neural induction media, differentiation media -VitA and differentiation media +VitA is as described before 21. For both experiments with YAP activator (Fig4, Extended Data Fig. 5), organoids were generated using the unlabelled WTC parent iPSC line. For the scRNAseq in Figure 4g-k, organoids were cultured as described above with addition of 1:1000 DMSO in neural induction medium to control organoids on day 5 and 10*µ*M Py-60 (MedChem Express, HY-141644) to organoids on day 5 or day 7. Media with fresh inhibitor or control media were exchanged every other day till dissociation and sequencing on day 10. For the scRNAseq in Extended Data Fig. 6, EBs were generated from 500 cells in mTSR+ (1:200 Rocki and 1:200 Pen-strep), per well of a 96 well plate and centrifuged at 200g for 5 min to generate embryoid bodies (EBs). Fresh mTSR+ with 1:200 ROCKi and 1:200 Pen-strep was exchanged on day 2 and day 4. Fresh Neural induction media was supplied on day 6 and exchanged every other day, followed by differentiation media -VitA on day 10 and differentiation media +VitA on Day 15. Organoids were given 2% matrigel, or 10*µ*M Py-60 with 2% matrigel or no matrix on day 10. All media compositions were exchanged on day 13 followed by dissociation and sequencing on day 16. The use of human ES cells for the generation of brain organoids was approved by the ethics committee of northwest and central Switzerland (2019-01016) and the Swiss federal office of public health.

#### Organoid dissociation and scRNAseq

For the time course data (Fig. 1), brain organoids were generated from the Histone2B-mEGFP cell line (cell line ID: AICS-0061-036). Organoids from day 5-11 belonged to the same batch, day 16-21 to a different batch and day 30 to a third batch. Multiple organoids of each line were pooled together to obtain a sufficient number of cells. For the early time points (Day 5,7,11), 24 organoids, each grown in an independent well of a 96 well plate were pooled, decreasing to 12 organoids for day 16, 10 organoids on day 21 and 6 on day 30. For the day 13 scRNAseq (Figure 4) with matrigel (11 organoids), no matrix (18 organoids) and agarose embedding (11 organoids), the organoids were generated from the unlabelled WTC parent iPSC line. Agarose was degraded using cell recovery solution (Corning, 11543560) at 4°C. For the experiments with YAP activator (Fig4), 16 control organoids, 23 organoids from Py-60 (given on day 5) and 20 organoids from Py-60 (given on day 7) were hashed together and used for scRNAseq. For the experiments in Extended Data Fig. 5, 5 control organoids (matrigel), 4 no matrix organoids and 13 organoids from Py-60 treatment were hashed and used for scRNAseq.

For all experiments, single-cell suspensions were generated by dissociation of the organoids with a papain-based neural dissociation kit (Miltenyi Biotec, 130-092-628). Briefly, organoids were washed three times with HBSS without Ca2+ and Mg2+ (STEMCELL Technologies, 37250). Prewarmed papain solution (1-2 ml) was added to the organoids and incubated for 15 min at 37°C. The tissue pieces were triturated 5–10 times with 1000 *µ*l wide-bore and then P1000 pipette tips. The tissue pieces were incubated twice for 10 min at 37°C with additional trituration steps in between and after with P200 and P1000 pipette tips. Cells were filtered consecutively with a 30 or 40*µ*m filter, centrifuged at 300g for 5 min and resuspended in cold PBS. The viability and cell count for the single cell suspensions were assessed using a Trypan Blue assay on the automated cell counter Countess (Thermo Fisher Scientific). Cell suspensions from day 5,7,11,16 and 21 were cryopreserved in Bambanker (Nippon genetics europe, BBH03) and stored at -20°C until the scRNA-seq experiments were performed. The cryopreserved single-cell suspensions of each time point were thawed by warming up the cryo for 1–2 min in a water bath at 37°C and directly centrifuged in 10 ml prewarmed DMEM with 10% FBS. Cells were washed twice with PBS + 0.04% BSA and filtered through a 40 *µ*m cell strainer (Flomi). For scRNA-seq, cells were resuspended to a final concentration after counting and viability checking that enabled targeting 8,000 cells and, in case the cell numbers were not sufficient, all cells were loaded. The scRNA-seq libraries were generated using the Chromium Single Cell 3’ V3 Library & Gel Bead Kit. Single-cell encapsulation and library preparation were performed according to the manufacturer’s protocol.

#### Light sheet microscopy

All cell lines used for imaging were procured from the Coriell Institute as described above. For imaging, EBs were embedded in 20-50% matrigel in neural-induction medium on Day 4, one organoid per microwell and up to 16 organoids in one sample chamber. After gelification of matrigel, neural induction medium was added to the sample chamber and exchanged every other day. For the imaging experiments with ECM perturbations in Figure 2 and 3, the sample mounting chamber has a vertical separation to segment it into 4 sub-chambers containing 4x organoids each. EBs (4) were either embedded in 20-50% matrigel dissolved in neural induction medium, or 4 EBs were covered with 0.6% low melting agarose and for the no matrix condition, 8 EBs were added to the sample chamber without any embedding. For the imaging shown in Extended Data Fig. 1g, EBs were aggregated from HES3 line (NKX2-1:GFP) using iPS Brew and rock inhibitor (1:200). After one day of aggregation, the EBs were transferred to a sample mounting chamber and embedded in matrigel. A neural patterning media (described in^40^) was employed for 14 days, complemented with SB432542 (Miltenyi, 130-106-543) and rh-Noggin (Miltenyi, 130-103-456) mediated dual SMAD inhibition from day 0 to day 9. After this, media was changed to neural differentiation media with vitamin A (composition previously described21) until day 21. The organoids were treated with the average morphogen concentration of SHH (140 ng/ml, Miltenyi, 130-095-727) from day 3-14 together with purmorphamine (0.21 *µ*m, Miltenyi, 130-104-465). 3.5 *µ*m XAV939 (Miltenyi, 130-106-539) was added from Day 0-9. Media was exchanged after every two days. Imaging was done with the LS1 Live light sheet microscope developed by Viventis Microscopy, using a 25x objective demagnified to 18.5x, with a field of view that was 710*µ*m and xy pixel size of 0.347*µ*m. Successive z steps were acquired every 2*µ*m for 201 steps. The frame rate for acquisition was 30 minutes for Fig 1g-i, Fig 2a-g and Fig 3b-f. The frame rate for acquisition was 60 minutes for Fig 2o-s, Fig 3g-n, Fig 4h and Extended Data Fig. 1g.

#### Fixation and Whole− mount HCR

For whole-mount staining, organoids were fixed overnight at 4°C on the nutator, washed 3-5 times in PBST, dehydrated with a PBST–methanol gradient (50%, 100%) and stored at − 20°C in 100% methanol until use. All probe sets were designed and provided by Molecular Instruments. The amplifiers and buffers were also ordered from Molecular Instruments (https://www.molecularinstruments.com/). HCR was performed according to the manufacturer’s protocol provided by Molecular Instruments with small changes. All 5 hairpins (B1-B5) conjugated with the following dyes were used per experiment Alexa-(488, Alexa-514, Alexa-545, Alexa-594, Alexa-639). Briefly, the samples were rehydrated with a series of graded methanol–PBST washes (25%, 50%, 75%, 100%) for 5 min each at 4°C on the nutator and washed an additional time with PBST. The samples were then treated with 10 *µ*g ml− 1 proteinase K (Invitrogen, 25530-049) for 3-5 min at room temperature followed by twice 2x PBST washes for 5 min. They were then post-fixed with 4% PFA for 20 min at room temperature and washed three times with PBST for 5 min each. The organoids were pre-hybridized in the probe hybridization buffer for 30 min at 37°C. 1 pmol of each probe set was diluted into probe hybridization buffer and the samples were incubated overnight at 37°C. Next day, the samples were washed 4x with the probe wash buffer at 37°C and washed 2x more with 5× SSCT. The organoids were then incubated in the amplification buffer for 10 min at room temperature followed by adding snap-cooled hairpin mixture diluted in the amplification buffer to incubate overnight at 25°C. The excess hairpins are washed the next day with 2x 5 min washes as well as 2x longer washes of 30 min followed by 1x 5 min wash with 5x SSCT buffer at room temperature. Organoids were stained with DAPI(1*µ*g/*µ*l) during the first 30 min washes. The samples were stored at 4°C and mounted on a *µ*-Slide chamber (Ibidi, 80807) and covered with 1% agarose. The samples were imaged using a 10x water immersion, 0.8 NA objective on the Zeiss LSM980 Airyscan system. Images were acquired using lambda scanning followed by spectral unmixing to image 6 channels in three sets of excitation (1st round 514 nm + 639 nm, 2nd round 488 nm + 545 nm + 594 nm, 3rd round 405nm). All images were processed using Fiji and the BigDataViewer plugin^41,42^.

#### Bulk CUT&Tag for YAP1

Organoids were generated from the WTC-11 iPSC line by culturing EBs (as described above) for 6 days, following which neural induction media was added on day 6. On day 10 differentiation media -VitA was given with 2% dissolved matrigel. Single cell suspensions of 12 days old organoids were prepared using the Miltenyi Neural Tissue Dissociation Kit (P) (130-092-628) following the manufacturers guidelines. Cells were counted and directly transferred into CUT&Tag Wash buffer supplemented with 0.01% Digitonin (20 mM HEPES pH 7.5; 150 mM NaCl; 0.5 mM Spermidine; 1× Roche Protease inhibitor cocktail). Per experiment, 1 Million cells were used and incubated with 2 *µ*g YAP antibodies (Abcam ab52771, Santa Cruz sc-101199). All following steps were performed as described^43,44^, except that the Tn5 incubation and cutting was performed at a NaCl concentration of 150 mM. To control unspecific cutting we performed the experiment without antibody (Tn5 only) and with a generic anti-rabbit antibody. The proteinA-Tn5 was purified in house following43. Final libraries were sequenced on the NovaSeq platform with PE 2×50 bp read length. Sequenced reads were mapped against hg38 using Bowtie2^45^. These were filtered for PCR duplicates and mapping quality. Coverage tracks were then generated using deeptools2 bamCoverage and normalized by sequencing depth^46^. Tracks were visualized using IGV.

#### Immunohistochemistry

For whole-mount staining, organoids were fixed overnight at 4°C on the nutator, washed 3 times in PBST, dehydrated with a PBST–ethanol gradient (50%, 100%) and stored at − 20°C in 100% ethanol until use. Before staining, organoids were dehydrated to 100% PBST and permeabilized+blocked in staining solution (0.5% Triton X-100, 0.2% Tween-20, 5% normal donkey serum, 1% Normal goat serum, 1% BSA in PBST) for 1 hour at room temperature. The samples were incubated overnight at 4°C in a staining solution containing rabbit anti-SOX10 (1:100, Abcam, ab227680).The next day, the samples were rinsed three times in PBST at room temperature and incubated overnight at 4°C with 1:500 secondary antibody (donkey anti-rabbit Alexa 568) in a staining solution. Samples were then washed 3-5 times with PBST for 5-10 minutes and stained with DAPI(1*µ*g/*µ*l) for 30 minutes followed by 2-3 PBST washes for 5 minutes each. The samples were stored at 4°C and mounted on a *µ*-Slide chamber (Ibidi, 80807) and covered with 1% agarose and immersed in PBST. The organoids were imaged using the Zeiss Airyscan confocal 980 microscope using a 10X water immersion objective. The images were further processed using Fiji. For cryosectioning, organoids were washed three times with PBST post fixation and then transferred to a 30% sucrose solution for 24–48 h at 4°for cryoprotection. Organoids were then transferred to plastic cryomolds (Tissue Tek) and embedded in OCT compound 4583 (Tissue Tek) for snap-freezing on dry ice. Organoids were sectioned in slices of 10 *µ*m thickness using a cryostat (Thermo Fisher Scientific, Cryostar NX50). The tissue slides were incubated in blocking-permeabilizing solution (0.3% Triton X-100, 0.2% Tween-20 and 5% normal donkey serum in PBS) for 1 h at room temperature. Next, the sections were incubated overnight at 4°C in a blocking-permeabilizing solution containing rabbit anti-SOX10 (1:100, Abcam, ab227680). The next day, the sections were rinsed three times in PBS before incubation for overnight at 4°C with 1:500 secondary antibody (donkey anti-rabbit Alexa 568) in blocking-permeabilizing solution. Finally, the secondary antibody solution was washed off with PBS and the sections were stained with DAPI before covering with ProLong Gold Antifade Mountant medium (Thermo Fisher Scientific). Stained organoid cryosections were imaged using the Zeiss Airyscan confocal 980 microscope using a 10X water immersion or 40X water immersion objective. For iterative histochemistry experiments, organoids were fixed at different timepoints and treated as described previously ^47^. The antibodies used were as follows, CDH2 (1:100, R&D Systems, AF6426), COL4A1 (1:25, Merck, AB769). The images were processed using Fiji.

### Data analysis methods

#### Preprocessing of scRNA-seq data from the organoid time course

We used Cell Ranger (v.3.0.2) with the default parameters to obtain transcript count matrices by aligning the sequencing reads to the human genome and transcriptome (hg38, provided by 10x Genomics, v.3.0.0). Count matrices were further preprocessed using the Seurat R package (v.4.3.0)^48^. First, cells were filtered on the basis of the number of detected genes and the fraction of mitochondrial genes. As sequencing depth varied between time points, the threshold of the number of detected genes was set individually for each sample. For the scRNAseq timecourse the number of detected genes is between 1000 to 7500 and the mitochondrial genes threshold is <10%. 3000 variable features were used and the number of PCA was set to 50. The total number of cells per day after preprocessing were (Day 5 = 5481, Day 7 = 8183, Day 7 = 4912, Day 16 = 6571, Day 21 = 7962, Day 30 = 7950).

#### Integration of different time points

Integration of time points was performed using the log-normalized gene expression data of meta cells. We used the union of the selected genes and TFs and further excluded cell-cycle-related genes from the set. Next, we computed cell cycle scores using the Seurat function CellCycleScoring(). Subsequently the data were z-scaled, cell cycle scores were regressed out (ScaleData()) and Principal Component Analysis (PCA) was performed using the Seurat function RunPCA(). We used the first 10 principal components (PCs) to integrate the different time points in the dataset using the CSS method. To remove any remaining cell cycle signal for any downstream tasks, we again regressed out the cell cycle scores from the integrated CSS matrix. To obtain a two-dimensional representation of the data, we performed UMAP embedding using RunUMAP() with spread =1.0, min.dist = 0.4 and otherwise the default parameters.

#### Pseudotime analysis of the early progenitor transition trajectory

Cells from day5, day7 and day 11 which were annotated as neuroectodermal (both N. Ect and Late N.Ect) were subsetted. CSS was used to re-integrate cells from different time points, and PCA was performed on the CSS representation to further reduce embedding dimensions to 10. A linear model was fit for each PCA-reduced CSS dimension, each with the cell cycle scores as independent variables. Residuals of the linear models were obtained as the cell-cycle-regressed-out PCA-reduced CSS representation, which was then used as the input of diffusion map (implemented in the destiny R package). The first diffusion component (DC1) was ranked across cells in the subset, and the ranked DC1 was used as the pseudotime representing the early progenitor transition progression.

#### Identification of genes with expression changed during early progenitor transition

For each gene detected in more than 1% of cells during early progenitor transition, two linear models were fit. The full model includes the natural spline (degree=5) of pseudotimes, while the reduced model only contains intercept. Two tests were performed to compare the two models. The relaxed test uses F-test to compare regression mean squares of the two models. The stringent test uses F-test to compare mean square errors of the two models. To correct for multiple testing, Bonferroni method was applied to the relaxed tests, and Benjamini-Hochberg method was applied to the stringent tests. Genes with adjusted P<0.05 in relaxed tests were defined as the broad set of genes with expression changed during the transition, while those with adjusted P<0.01 in stringent tests were defined as the stringent set.

#### Preprocessing of scRNA-seq data from all other datasets

We used Cell Ranger (v.3.0.2) with the default parameters to obtain transcript count matrices by aligning the sequencing reads to the human genome and transcriptome (hg38, provided by 10x Genomics, v.3.0.0). Count matrices were further preprocessed using the Seurat R package (v.4.3.0). The cells were filtered on the basis of the number of detected genes and the fraction of mitochondrial genes. As sequencing depth varied between time points, the threshold of the number of detected genes was set individually for each sample. For the matrix perturbation dataset (Figure 4), the number of detected genes is between 500 to 8000, 3000 variable features were used and the number of PCA was set to 50. Mitochondrial and histone related genes were removed followed by integration. The total number of cells per condition after pre-processing were (Matrigel = 8095, no matrix = 8343, agarose = 3725). To obtain a two-dimensional representation of the data, we performed UMAP embedding using RunUMAP() with the default parameters. For the YAP perturbation (Figure 4), the number of detected genes is >200 and mitochondrial genes threshold was set to <20%. 3000 variable features were used and the number of PCA was set to 50. Mitochondrial and histone related genes were removed followed by integration. The total number of cells per condition after pre-processing were (Control = 2085, Py-60 given on day 5 = 763, Py-60 given on day 7 = 1955). To obtain a two-dimensional representation of the data, we performed UMAP embedding using RunUMAP() with spread =0.3, min.dist = 0.5. For the scRNAseq datasets in Extended Data Fig. 6, the number of detected genes >1000 and percent.mt < 10. 3000 variable features were used and the number of PCA was set to 20. The total number of cells per condition after pre-processing were (Matrigel = 1183, no matrix = 1285, YAP activator Py-60 = 1189). To obtain a two-dimensional representation of the data, we performed UMAP embedding using RunUMAP() with the default parameters. For all datasets, transcript counts were normalized to the total number of counts for that cell, multiplied by a scaling factor of 10,000 and subsequently natural-log transformed (NormalizeData()). The datasets in different experiments were integrated using CSS: cluster similarity spectrum ^49^.

### Light sheet image analysis

#### Denoising and background subtraction

To increase the comparability between images in the dataset all movies were denoised using Noise2void^50^ (v0.3.1). A separate 2D denoiser was trained for every position and every channel. The denoiser was trained on randomly selected z-slices from throughout each corresponding movie and channel. For the datasets used in Fig. 1g-i, Fig. 2a-g, Fig. 3b-f and Extended Data Fig. 1f 30 randomly selected z-slices were used and for the dataset in Fig. 2o-s and Fig. 3g-n 90 random-z slices were used. The randomly selected z-slices were split into a training subset with 90 percent of the z-slices and a test subset with 10 percent of the z-slices. The denoiser was trained for 128 epochs on the training subset with 96 by 96 pixel patches, with a batch size of 256, a unet_kern_size of 3, train_loss equal to mse, n2v_perc_pix of 0.198. The training progress was monitored on the test subset. The trained denoiser was then used to denoise all images in the movies. After this, the background subtraction was performed by using the minimal value along the z-axis and clipping small intensity values. The Extended Data Fig.1f dataset was further dehazed using the dehazing function from dexp^51^ (v2022.6.29.552) with a filter size of 60 and correct_max_level set to false. For Fig. 1i and for Fig. 2o the 3D image were trilinearly rescaled to an isotropic voxel size of 1.15 *µ*m. Then a wedge mask was created with a triangular cutout and then multiplied with the downscaled image. Then the image was rotated using scipy^52^ (v1.7.3) ndi.rotate function. The images were then attenuated as implemented in the dexp.processing.color.projection function with the parameters attenuation of 0.0005, attenuation_min_density of 0.3 and attenuation_filtering of 24. For the spinning movie in Supplementary video 2 the 3D image was trilinearly rescaled, with antialiasing, to an isotropic voxel size of 1.388 *µ*m and 32 zero padding planes have been added to before and after in x,y and z directions. The image was then rotated with the scipy.ndi rotate function and after each rotation each channel is attenuated with the parameters attenuation_filtering=4, attenuation_min_density= 0.002 and attenuation=0.01.

#### Temporal registration

The light sheet dataset from Fig. 1g-i, Fig. 2a-g, Fig. 3b-f was registered. First, the movies were cropped and centered using the cropping algorithm implemented in LStree^53^ (v0.1). As only one organoid is present per position, the largest object was linked throughout the movie for cropping. After cropping the images were padded with zeros, 32 planes before and after the image in the z-direction, and with 96 planes before and after the image in both x and y direction prior to registration. For the temporal image registration, translational registration from itk-elastixs^54,55^ (v5.2.1) with the default translation parameter map was used to register the movies using GFP as the leading channel. The first frame in the movie was used as a reference frame and then the second image was registered to it. After this the translation was applied to both channels of the second image. Then the next image was registered to the previously registered frame in the movie and again the translation was applied to both channels in the next image. This was repeated until all images in a movie were registered. After translation, the images were clipped to ensure that no small negative values remained. Movies in the positions 9 to 16 contained some higher intensity areas. To deal with this, prior to calculating the registration, the image intensities were rescaled to the first and 99th percentile and the default translation parameter map with three resolutions was used for registration. The calculated translation was then applied to both non intensity rescaled images for these positions.

#### Tissue segmentation

A pixel wise random forest classifier was used for the tissue segmentation similar to other already established tools such as Weka segmentation^56^. Due to the multi-terabyte size of the dataset a python pixel wise segmentation classifier using both scikit-learn^57^ (v0.18.3) and scikit-image^58^ (v1.1.1) was implemented (https://scikit-image.org/docs/stable/auto_examples/segmentation/plot_trainable_segmentation.html#id4). The input for the random forest classifier were 2D image features from the sum of both channels, which were extracted using multiscale basic features function from scikit-image with a minimum sigma of 1, a maximum sigma of 64, intensity, edge and texture features. The input image was bilinearly downscaled to a quarter resolution in xy. To train the classifier, random z-slices from throughout the movies were annotated with bounding boxes in jupyter notebook using bboxwidgets (v0.4.0)(https://github.com/gereleth/jupyter-bbox-widget), as either background, lumen or organoid tissue. Random slices were iteratively annotated until satisfactory image segmentations were produced. In total 1414 slices were corrected/annotated for the dataset used in Fig. 2c-f, Extended Data Fig. 3a-c. and 2221 slices for the dataset used in Fig. 2p-s and Extended Data Fig. 3d,e. For every slice with a bounding box the 2D image features were extracted and a random forest classifier was trained to predict the labels. One classifier was trained for each dataset. The random forest classifiers for the two datasets were trained to a depth of 25, with 50 random trees and max_samples of 0.05. The trained models were then used to predict the labels for each pixel for each z-slice in each time point of the movies. For both datasets the segmentation masks were predicted for one time point per hour. After predicting the labels for each 2D segmentation, the output masks were stacked to a 3D segmentation mask. After this the organoid and the lumen masks were extracted and smoothed by applying a scikit-images gaussian filter with a voxel size of 1.388 in the z-direction and 2 in both the the x and y directions to the binary mask and then rebinarizing and combining the masks back to one segmentation image. After this step each z-slice of the 3D segmentation masks were further processed in 2D. First small lumen were removed using the binary_closing function from scikit-image with a disk size of 3. Any holes in the lumen masks were filled and any background pixels within the organoid were assigned the lumen label. Further, any voxels labeled lumen outside of the convex hull of the organoid were assigned the background label. The slice wise 2D segmentations were then reassembled into a 3D tissue segmentation. As only one organoid is present in one movie, label and regionprops functions from scikit-image were used to extract the largest non-background object for further downstream analysis. The resulting masks were visualized in 3D for Fig. 2c,p, Extended Data Fig. 3b,d, Supplementary video 5 and Supplementary video 9. The masks were rescaled to an isotropic resolution of 1.388 *µ*m, extracting each lumen surface using scikit-images marching cubes, with a step_size of 1 and level 0 and the organoid surface using marching cubes with a step_size of 2 and level 0. Next the surface meshes were cleaned using pymeshfix^59^ (v0.16.1) (https://github.com/pyvista/pymeshfix/tree/main) clean_from_arrays and finally the meshes were smoothed using the filter_taubin function from trimesh^60^ (v3.13.0) (https://github.com/mikedh/trimesh) with 50 iterations. The surface meshes were then plotted using matplotlib (v3.5.2) plot_trisurf with the bounding box being equal to the maximum length in any of the xy and z direction in *µ*m. For Extended Data Fig. 3a, Supplementary video 4 and Supplementary video 8, using the organoid masks the 90th percentile largest organoid z-slice was calculated for every time point in every movie. Then a sliding average was applied over the calculated 90th percentile z-slices with a window size of 20 for Extended Data Fig. 3a and Supplementary video 4 and a window size of 5 for Supplementary video 8. Lastly for each of these slices, the images were false colored according to the masks.

#### Tissue quantification

To characterize the tissue architecture, the organoid tissue and lumen sizes were quantified for the first 124 hours for both Fig. 2d-f and Fig. 2q-s. For the lumen quantification, any lumen with a size smaller than 20’000 *µ*m^3^ were removed using scikit-images remove_small_objects function. Further scikit-images label and regionprops functions were used to assess the number of lumen and the size of each lumen. For Fig. 2d-f and Extended Data Fig. 3c for the organoid at position 2 the measurements were not considered after hour 49, as the camera moved to another position and for the organoid at position 4 between hours 43 to 48 were not considered as the organoid moves out of the field of view. For the measurements in Fig. 2q-s and the Extended Data Fig. 3e in addition to the 4 Matrigel organoids already present in the experiment, 4 randomly picked organoids from the experiment in Fig. 2d-f were used to supplement the measurements.

#### Single cell segmentation

To quantify changes in cell morphologies, 3D cell segmentation was performed using EmbedSeg^26^ (v0.2.5). One 3D image for every 24 hours, for each movie was segmented. For the dataset in Fig. 3b-f, Extended Data Fig. 4b-f one position was segmented, for Fig. 3g-n Extended Data Fig. 5a-d 3 positions per condition were segmented. The Agarose condition was further supplemented with one additional red channel movie. Prior to training the model the images were bilinearly downscaled by a factor of 2 in the xy direction. Hand selected 3D volumes were annotated using labkit^61^ and then used to train the model in a human in the loop fashion, where iteratively predicted 3D volumes are corrected and then used to retrain the model. In total 5850 cells were hand corrected in this fashion. For creating the crops for training the model, the medoid was used and a n-sigma of 4 used to calculate the crop_size. Further a speed_up factor of 3 was used to create the crops. The model tile size was set to 600 for both the x and y directions and 80 in the z-direction. The model was trained on 5512 cells and 338 cells were set aside as a validation set. The 3D model was trained for 200 epochs, with a batch size of 8 and then the model which had the best iou performance of 0.72 on the validation set was selected. Using tiling the 3D volumes were segmented, with a fg_thresh of 0.4 and seed_thresh of 0.7 and test-time-augmentation set to True.

#### Cell morphology quantification

Morphometrics^62^ (v0.0.6)(https://github.com/morphometrics/morphometrics) was used to extract cell morphology measurements from the segmented images. The images were rescaled trilinearly and the masks with a nearest neighbor interpolation to an isotropic voxel size of 0.694 *µ*m. To speed up the measurements, masks with a volume smaller than 100 voxels (33.4 *µ*m^3^) were removed using remove_small_objects from scikit-image. The surface, size, intensity, position and moments morphometric measurements were extracted for each cell mask using morphometrics. Additionally the major and the minor axis length for each cell was measured using the scikit-images regionprops function.

#### Quality control and demultiplexing

Prior to the manual quality control steps, masks with a volume of lower than 100 *µ*m^3^ and a maximum intensity of less than 20 were removed from further analysis leading to a total of 299’027 subcellular structures. Further, any measurements containing all NaNs and after this cells containing any NaNs were also removed from the analysis. Scanpy^63^ (v1.9.1) and morphometrics were used to normalize, dimensionally reduce using PCA and cluster (Leiden) the dataset. Then random cells were extracted, from different time points, positions and clusters and labeled to either be a Histone, Lamin, Actin, Tubulin, cell membrane or a faulty mask. In total 3379 cells were hand annotated in this fashion. The annotated cells were then split into a train (90%) and a test set (10%). For demultiplexing all measured measurements from morphometrics and the major and minor axis were used. The training data was used to perform a 4-fold cross validation grid search on random forest classifiers from scikit-learn on the number estimators and the maximum depth of the tree. The best performing random forest classifier was selected for demultiplexing and quality control. The model achieved an accuracy of 0.73 on the test set. After this, wrongly demultiplexed structures, which either did not exist in the experiment or originated from the wrong channel were removed. After manual quality control a total of 59’053 structures remained.

#### Morphology analysis

All morphology analyses were run using surface, size and axis length measurements and additionally the axis length ratio was added. Scanpy^63^ was used to morphologically analyze the segment markers. For Fig. 3g-k,n only Actin cells from the ECM perturbation experiment were used and the measurements were scaled and dimensionally reduced using the pca function from morphometrics. The neighborhood graph was then calculated using the first five principal components and then clustered using Leiden clustering with a resolution of 0.6 using the cluster_features function from morphometrics. After this, clusters 8 and 9 contained only debris and were removed from the further analysis and the data was rescaled, dimensionally reduced and clustered with the same parameters. After clustering a graph embedding was calculated using PAGA^28^ with a threshold of 0.1 and the daily proportions of the different clusters for the conditions were calculated. For Fig. 3g,h, to obtain a two dimensional representation of the data, a PAGA initialized UMAP embedding was calculated. Based on the axis length ratio of the cells in the clusters, the clusters were either colored as containing elongated (cyan) or non-elongated cells (yellow). To create Fig. 3j, the first non-zero element from the 3D Actin masks used in Fig. 3g-k was extracted. This was then used to extract a 2D image with the corresponding voxels from the intensity image and to color the pixels according to the cluster membership. To estimate the diversity of the cell morphologies throughout the time series, the Shannon index was calculated for every time point and every condition: 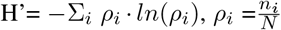 where N is the total number of cells of all clusters per day and condition, and *n*_*i*_ is the number of members of a cluster for a specific condition and day. For Fig. 3d,e all demultiplexed markers were used and the neighborhood graph was then calculated using the first 4 principal components and then clustered using Leiden clustering with a resolution of 1.5. After clustering a graph embedding was calculated using PAGA^28^ with a threshold of 0.1 and a PAGA initialized UMAP embedding was calculated. For Extended Data Fig. 4b-f and Extended Data Fig. 5b the neighborhood graphs were then calculated using the first 4 principal components and then clustered using Leiden clustering with a resolution of 0.4. After this the PAGA graphs and the PAGA initialized UMAPs were calculated.

#### Cell angle analysis to the surface

To calculate the cell angle to the surface, first an ellipsoid was used to fit to the surface of the 3D masks of the individual cells (https://github.com/aleksandrbazhin/ellipsoid_fit_python). The surface of the binary subcellular feature masks were extracted using marching_cubes function from scikit-image with a step size of 2 and level 0. Then, to estimate the directionality of the cell masks, the major axis of the 3D ellipsoid was extracted. Any cells where the ellipsoid had a negative radii or a radii larger than 180 voxels were excluded from further analysis. The organoid mask was upscaled to the same dimensions as the cell segmentation masks and used to extract the nearest surface normal for each cell. To obtain the surface normals, a surface mesh was extracted using scikit-images marching cubes, with a step_size of 40 and level 0. For all surface meshes pymeshfix clean_from_arrays function was used to clean the surface mesh. For the organoid surface mesh, the filter_taubin function from trimesh with 50 iterations was used to smoothen the surface. spatial.cKDTree from scipy (v.1.8.1) was used to obtain the nearest surface normal from the centroid of each cell. We calculated the alignment index as the absolute cosine similarity (scikit-image) between the major axis of the ellipsoid and the nearest surface normal of the organoid. For Fig. 3f,m,l and Extended Data Fig. 5c only Actin was considered, while for Extended Data Fig. 5d all demultiplexed markers were used. All plots for Fig. 3f,l and for Extended Data Fig. 5d were created using the ellipsoid_plot from (https://github.com/aleksandrbazhin/ellipsoid_fit_python).

## Extended Data Figures

**Extended Data Fig. 1.**
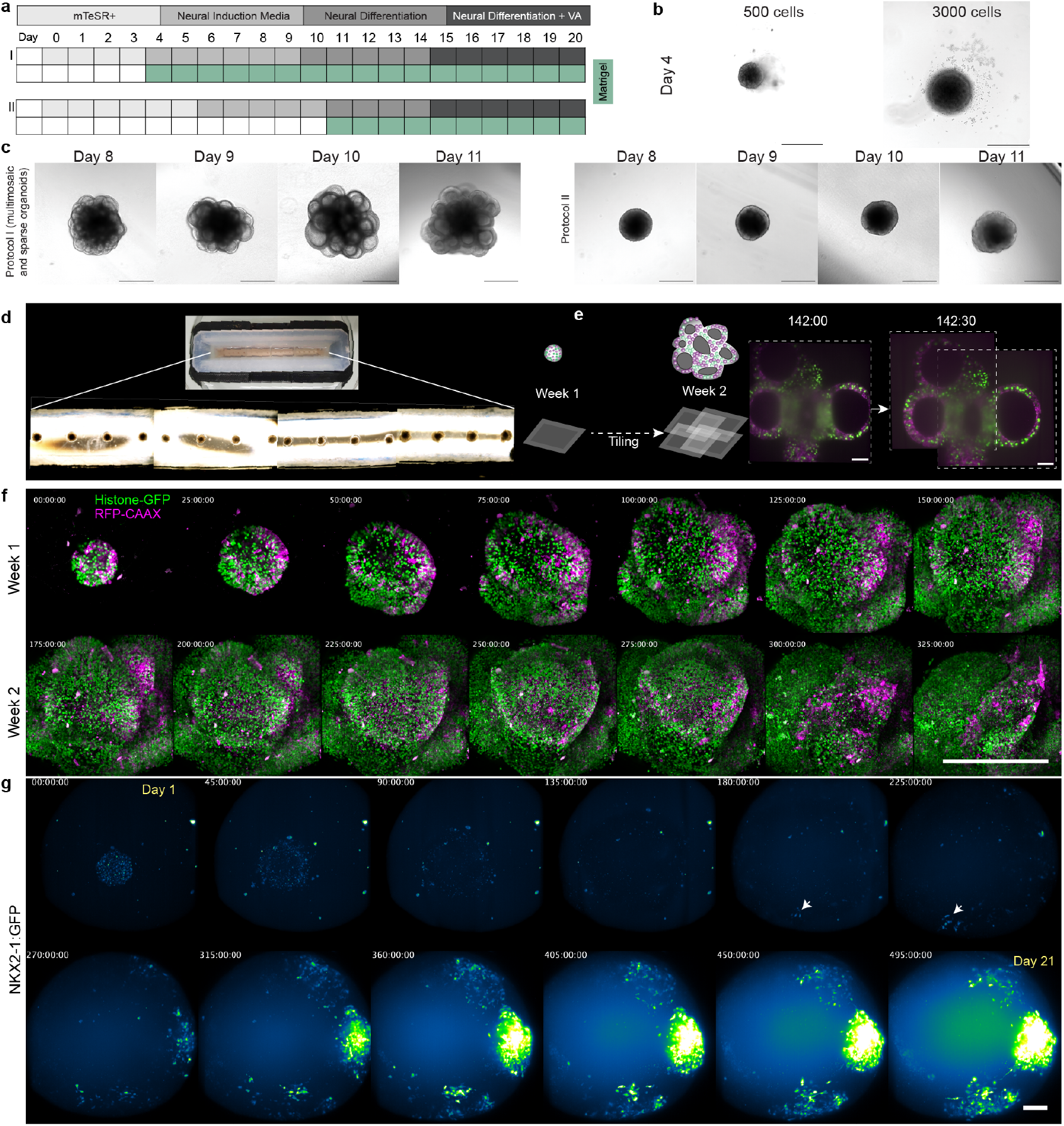
Long-term live imaging of fluorescent brain organoids. a) Schematic of the protocol developed in this study (Protocol I) and a previous protocol to develop multi-region brain organoids (Cerebral organoid, Protocol II)1,21. b) Comparison of organoid size at day 4 post aggregation at two cell seeding concentrations. Scale Bar is 500 micrometers. c) Brightfield images showing organoid growth from day 8-11 in the two different protocols.Scale Bar is 500 micrometers. d) Photographs of the sample holding chamber with four separated sub-chambers each containing four microwells, for growing and imaging organoids with sixteen different organoids arranged one per microwell. e) Schematic overview of tiled image acquisition and example images showing two timepoints, before and after tiled acquisition and image fusion. Scale Bar is 100 micrometers.Time is in hours. f) Fluorescent image stills from a 2 week imaging acquisition of a mosaic organoid (Histone2B-GFP, green; RFP-CAAX, pink, unlabelled WTC-11).Scale Bar is 500 micrometers. Time is in hours. g) Images from a 3-week continuous imaging experiment using a NKX2-1:GFP reporter line. The organoids were given SHH morphogen treatment to induce ventral telencephalic patterning of the organoids. Images are false-colored with the green-fire-blue LUT. Scale Bar is 100 micrometers. Time is in hours.

**Extended Data Fig. 2.**
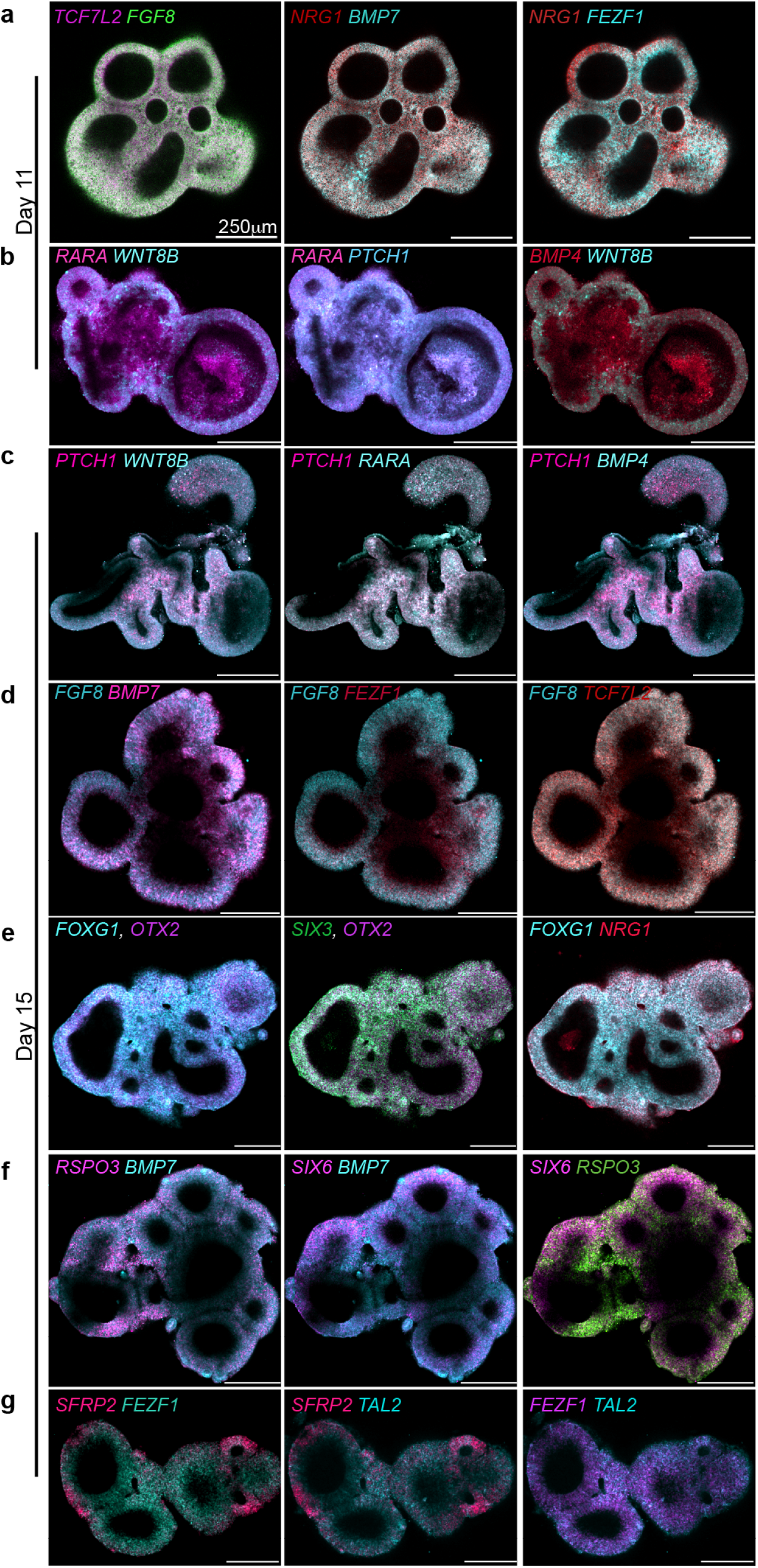
Fluorescence *in situ* staining (HCR) regional marker genes. Cross-section images of fluorescence *in situ* hybridization chain reaction (HCR) stainings showing RNA expression of marker genes in organoids fixed on day 11 (a-b) and day 15 (c-g). Scale, 250 micrometer in all images. Organoids were stained whole-mount and images were acquired as z-stacks. Selected cross-sections after reorientation of the acquired volume are shown for each organoid.

**Extended Data Fig. 3.**
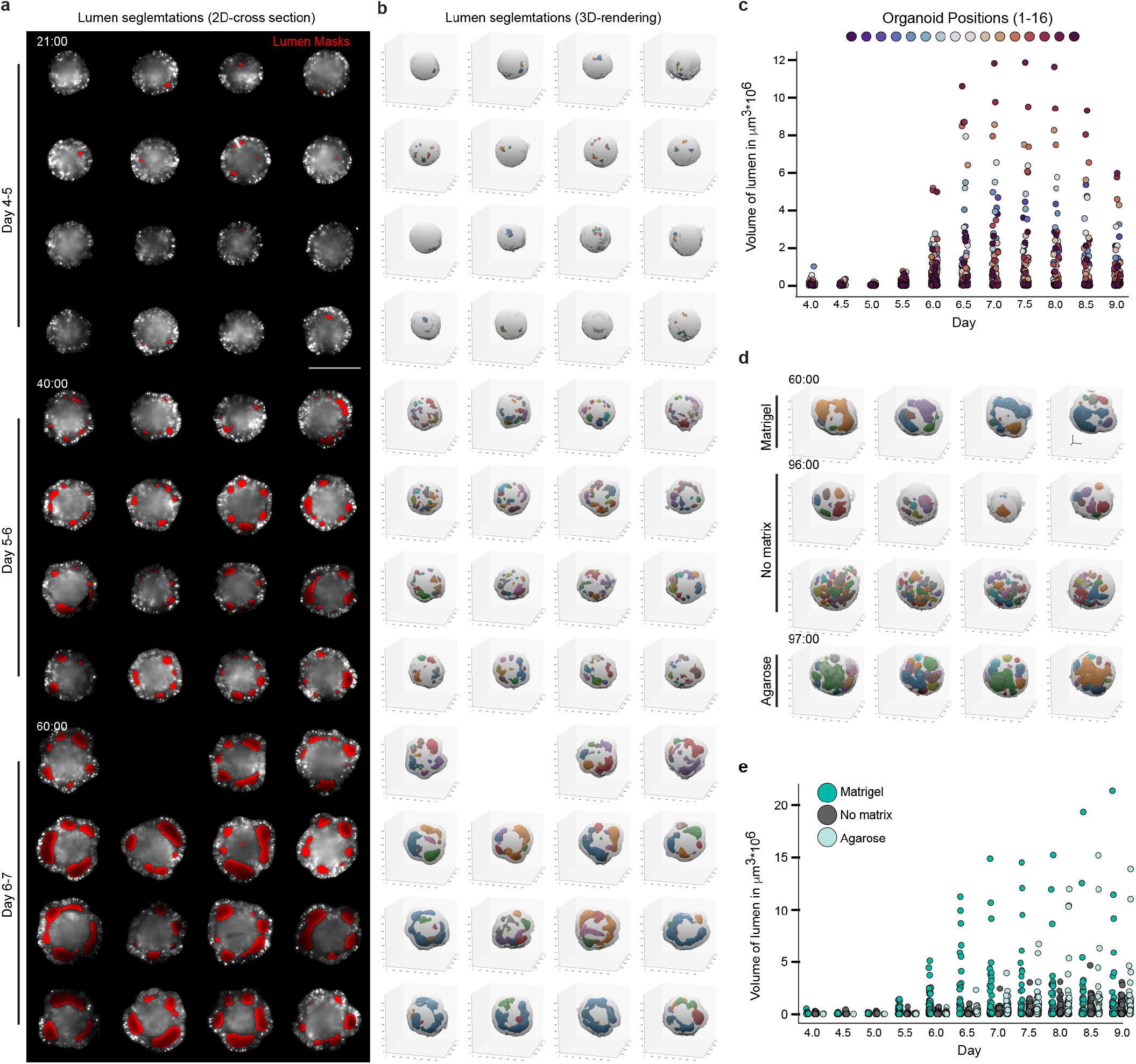
Lumen segmentations to track organoid and lumen morphodynamics in organoid development. a) 2D cross sections showing organoid epithelium (gray) and segmented lumen (red) from day 4-7. Scale, 500 micrometers. Time is in hours. All 16 organoids were imaged in one experiment. b) 3D renderings of segmented lumen for the corresponding organoids shown in a. Time is in hours. c) Plot shows individual lumen volumes over time, from day 4 to day 9, of all segmented lumen from organoids shown in a and b. d) 3D renderings of lumen segmentions in 4 different organoids in matrigel condition, 8 organoids from no matrix condition and 4 organoids for diffusion barrier condition. All 16 organoids were imaged in one experiment. Time is in hours. e) Plot shows individual lumen volumes from day 4 to day 9 of all segmented lumen in organoids treated with three different conditions (Matrigel (n=8), no matrix (n=8) and Diffusion barrier (n=4)). The combined measurements for matrigel conditions includes 4 random organoids picked from the experiment shown in c.

**Extended Data Fig. 4.**
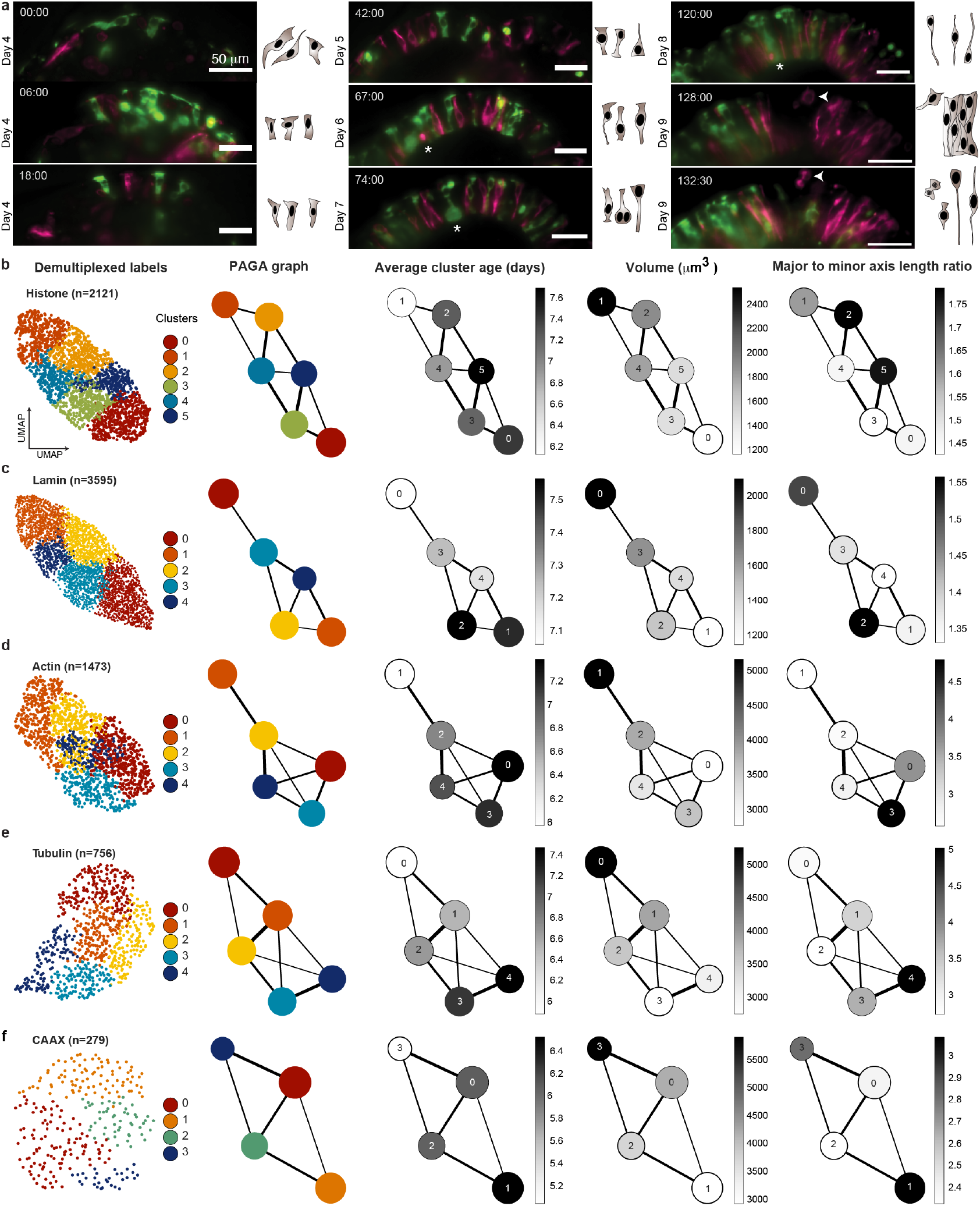
Demultiplexed labels reveal cellular morphodynamics. a) Cross-sections showing cell shape transitions from embryoid bodies to different stages of neuroepithelium formation and maturation in organoids from day 4 to day 9. Illustrations show different cell shapes seen in the cross-section images. Organoids are labeled with nuclear membrane (Lamin, RFP, magenta), plasma membrane label (CAAX, RFP, magenta), Actin (GFP, green), Tubulin (RFP, magenta), and nuclei (Histone, GFP, green). Asterisks mark cell division events seen in Tubulin (magenta) and Actin(green) labels. Arrowheads mark cell divisions on the basal surface of the organoids. Time is in hours. Scale, 50 micrometers. b-f) UMAP embeddings and PAGA plots showing cell morphotype clusters of all 5 demultiplexed labels based on morphometric feature extraction. The PAGA plots per label show average cluster age, the change in nuclei volume and the change in axis length measured over time across all segmented cells. n number indicates the total number of cells per label from all timepoints.

**Extended Data Fig. 5.**
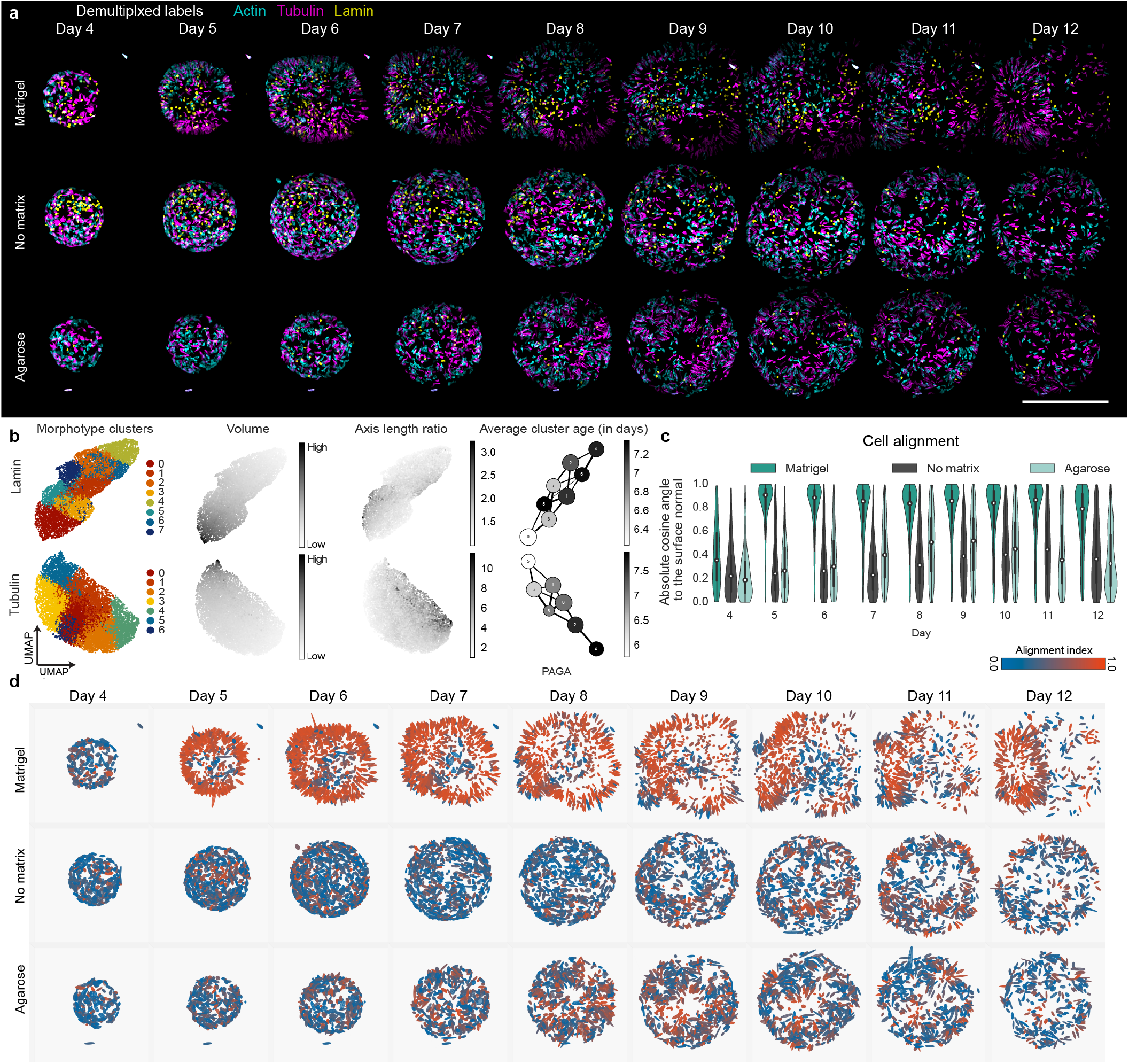
Demultiplexed labels reveal cellular morphodynamics in matrix perturbations. a) Demultiplexed images (Lamin, yellow; Actin, cyan; Tubulin, magenta) of example organoids that were embedded with matrigel, given no matrix or embedded in agarose and imaged from day 4-12 in a light sheet microscope. Scale, 500 micrometer. b) UMAP embeddings of Lamin and Tubulin labels based on morphometric feature extraction. UMAP embeddings show change in axis length and volume measured using cell segmentations. PAGA plots show change in average cluster age (days). c) Violin plot showing the cell alignment values across all segmented cells (Actin) from day 4 to day 12 for all three matrix conditions. d) Images show cells colored by their alignment index (absolute cosine angle to the nearest organoid surface normal) in all three conditions over time. Red colors indicate higher alignment (perpendicularity) to the cell surface.

**Extended Data Fig. 6.**
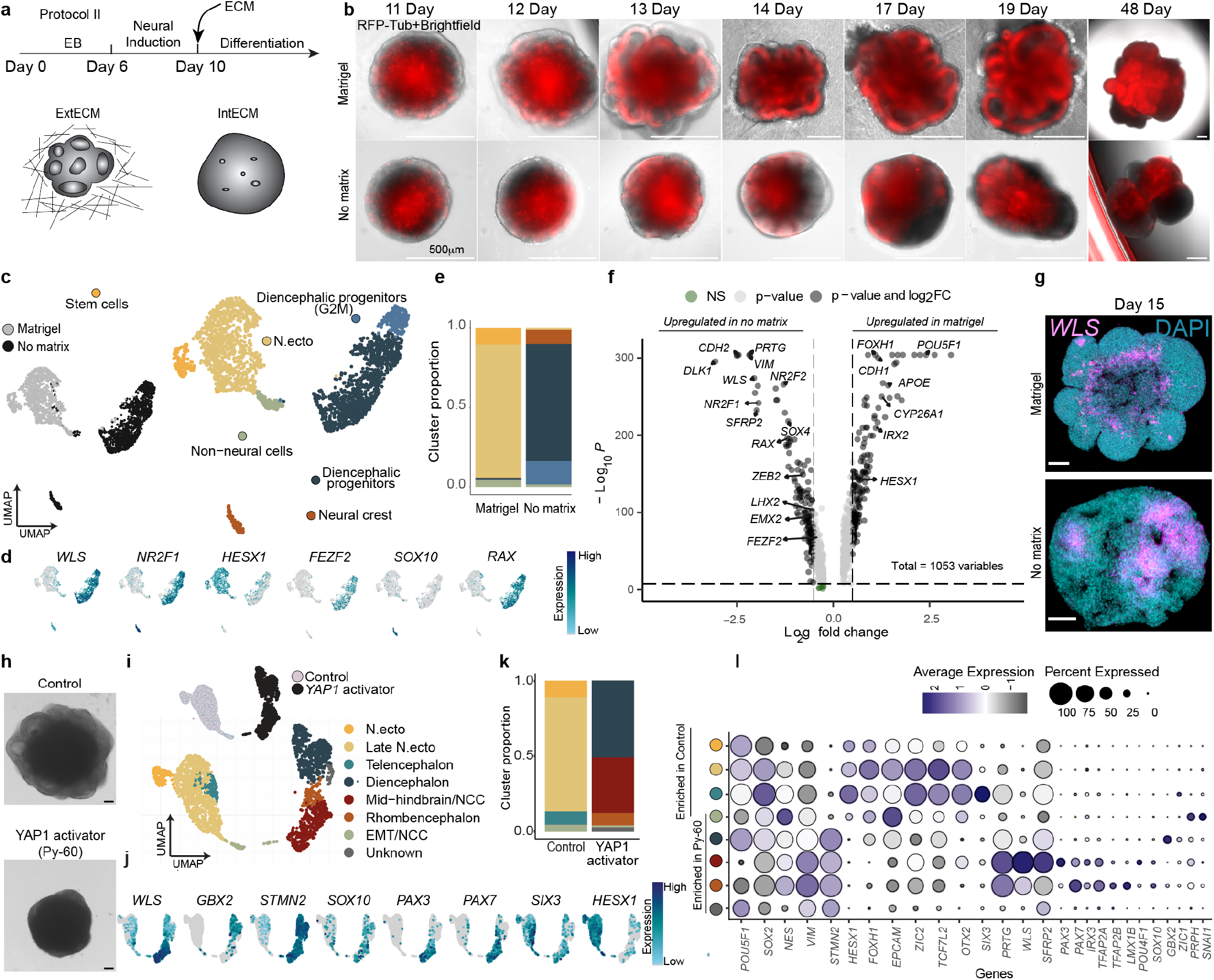
Matrigel influences organoid regional patterning. a) Overview of the protocol used to generate organoids to test the influence of matrix on brain organoids development (Cerebral organoid, Protocol II). b) Time-course images of organoids labeled with RFP-Tubulin showing brightfield and RFP expression in the presence and absence of matrigel. Imaging was done with a widefield Nikon Ti2 microscope. c) UMAP embeddings of scRNA-seq data performed at day 16 with cells labeled by treatment (left) or colored by cluster and labeled by cell population (right). d) Feature plots showing normalized expression of selected marker genes. e) barplot showing cluster proportions of the cell populations in control and YAP1 activator. f) Volcano plot showing differentially expressed genes upregulated in matrigel (right) and upregulated in no matrix conditions (left). g) Images show maximum intensity projections of organoids treated with matrigel (top) and grow in with no matrix (bottom) from a whole-mount fluorescence *in situ* hybridization chain reaction (HCR) staining at day 15. Scale, 100 micrometer. h) Brightfield images at day 13, showing a control organoid and an organoid cultured with YAP1 activator together with matrigel from day 10 onwards. Organoid protocol is shown in a. Scale, 100 micrometer. Organoids were used for an scRNAseq experiment on day 16 (i-l). i) UMAP embedding of scRNA-seq data from 16 day old organoids with cells colored by treatment (top) and cluster and labeled by cell population (bottom). N. ecto is neurectoderm, NCC is neural crest cells and EMT is epithelial-to-mesenchymal transition. j) Feature Plots showing normalized expression of marker genes. k) Stacked barplot showing the proportion of each cell population in control and YAP1 activator treatment. l) Dotplot showing average expression and percentage of cells expressing selected regional marker genes used for annotating the cell populations shown in i.

**Extended Data Fig. 7.**
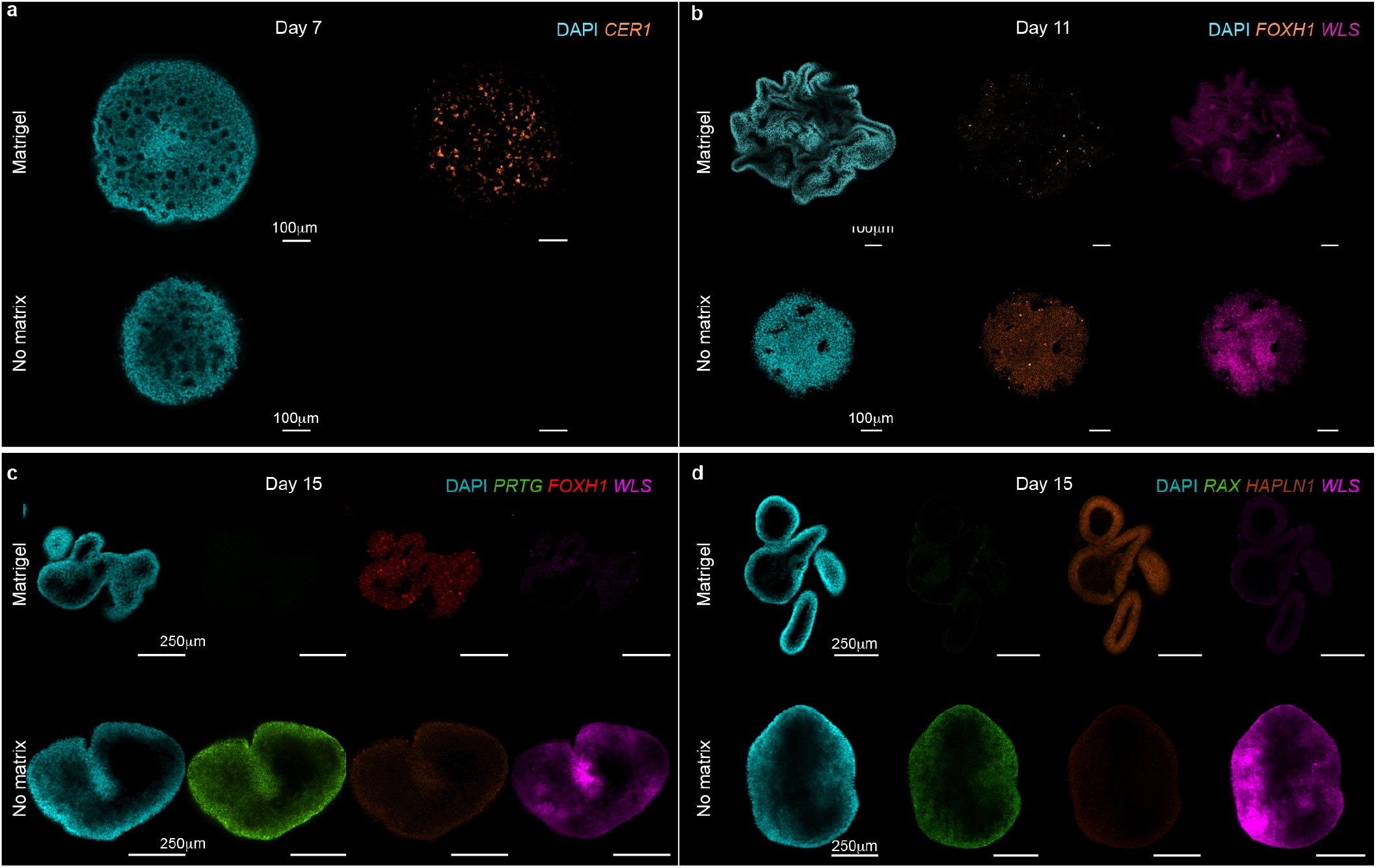
HCR stainings in organoids cultured with matrigel and with no extrinsic matrix. a) Cross section images of fluorescence *in situ* hybridization chain reaction (HCR) stainings showing RNA expression of marker genes between organoids grown with matrigel or with no matrix and fixed on day 7 (a), day 11 (b) and on day 15 (c-d). Scale, 250 micrometer in all images. a) Organoids show higher expression of Cer1 in matrigel day 7. b) Organoids show higher expression of FOXH1 in matrigel and of WLS in no matrix. c) Organoids show higher expression of FOXH1 in matrigel and of PRTG and WLS in no matrix. d) Organoids show higher expression of HAPLN1 in matrigel and of RAX and WLS in no matrix.

## REFERENCES

1. Lancaster, M. A. et al. Cerebral organoids model human brain development and microcephaly. Nature 501, 373–379 (2013).

2. Camp, J. G. et al. Human cerebral organoids recapitulate gene expression programs of fetal neocortex development. Proc. Natl. Acad. Sci. U. S. A. 112, 15672–15677 (2015).

3. Fleck, J. S. et al. Inferring and perturbing cell fate regulomes in human brain organoids. Nature 1–8 (2022).

4. Sasai, Y. Cytosystems dynamics in self-organization of tissue architecture. Nature 493, 318–326 (2013).

5. Renner, M. et al. Self-organized developmental patterning and differentiation in cerebral organoids. EMBO J. 36, 1316–1329 (2017).

6. Eiraku, M. & Sasai, Y. Mouse embryonic stem cell culture for generation of threedimensional retinal and cortical tissues. Nat. Protoc. 7, 69–79 (2011).

7. Eiraku, M. & Sasai, Y. Self-formation of layered neural structures in threedimensional culture of ES cells. Curr. Opin. Neurobiol. 22, 768–777 (2012).

8. Kanton, S. et al. Organoid single-cell genomic atlas uncovers human-specific features of brain development. Nature 574, 418–422 (2019).

9. Long, K. R. & Huttner, W. B. The Role of the Extracellular Matrix in Neural Progenitor Cell Proliferation and Cortical Folding During Human Neocortex Development. Front. Cell. Neurosci. 15, 804649 (2022).

10. Long, K. R. et al. Extracellular Matrix Components HAPLN1, Lumican, and Collagen I Cause Hyaluronic Acid-Dependent Folding of the Developing Human Neocortex. Neuron 99, 702–719.e6 (2018).

11. Fietz, S. A. et al. Transcriptomes of germinal zones of human and mouse fetal neocortex suggest a role of extracellular matrix in progenitor self-renewal. Proc. Natl. Acad. Sci. U. S. A. 109, 11836–11841 (2012).

12. Zhang, Z., O’Laughlin, R., Song, H. & Ming, G.-L. Patterning of brain organoids derived from human pluripotent stem cells. Curr. Opin. Neurobiol. 74, 102536 (2022).

13. Kelley, K. W. & Pas, ca, S. P. Human brain organogenesis: Toward a cellular understanding of development and disease. Cell 185, 42–61 (2022).

14. Sidhaye, J. & Knoblich, J. A. Brain organoids: an ensemble of bioassays to investigate human neurodevelopment and disease. Cell Death Differ. 28, 52–67 (2021).

15. Velasco, S. et al. Individual brain organoids reproducibly form cell diversity of the human cerebral cortex. Nature 570, 523–527 (2019).

16. Paşca, A. M. et al. Functional cortical neurons and astrocytes from human pluripotent stem cells in 3D culture. Nat. Methods 12, 671–678 (2015).

17. Serra, D. et al. Self-organization and symmetry breaking in intestinal organoid development. Nature 569, 66–72 (2019).

18. Karzbrun, E. et al. Human neural tube morphogenesis in vitro by geometric constraints. Nature 599, 268–272 (2021).

19. He, Z. et al. Lineage recording in human cerebral organoids. Nat. Methods 19, 90–99 (2022).

20. Viana, M. P. et al. Integrated intracellular organization and its variations in human iPS cells. Nature 613, 345–354 (2023).

21. Lancaster, M. A. & Knoblich, J. A. Generation of cerebral organoids from human pluripotent stem cells. Nat. Protoc. 9, 2329–2340 (2014).

22. Roberts, B. et al. Systematic gene tagging using CRISPR/Cas9 in human stem cells to illuminate cell organization. Mol. Biol. Cell 28, 2854–2874 (2017).

23. Long, K. R. & Huttner, W. B. How the extracellular matrix shapes neural development. Open Biol. 9, 180216 (2019).

24. Ethell, I. M. & Ethell, D. W. Matrix metalloproteinases in brain development and remodeling: synaptic functions and targets. J. Neurosci. Res. 85, 2813–2823 (2007).

25. Pluen, A., Netti, P. A., Jain, R. K. & Berk, D. A. Diffusion of macromolecules in agarose gels: comparison of linear and globular configurations. Biophys. J. 77, 542–552 (1999).

26. EmbedSeg: Embedding-based Instance Segmentation for Biomedical Microscopy Data. Med. Image Anal. 81, 102523 (2022).

27. Becht, E. et al. Dimensionality reduction for visualizing single-cell data using UMAP. Nat. Biotechnol. (2018) doi:10.1038/nbt.4314.

28. Wolf, F. A. et al. PAGA: graph abstraction reconciles clustering with trajectory inference through a topology preserving map of single cells. Genome Biol. 20, 59 (2019).

29. Braun, E. et al. Comprehensive cell atlas of the first-trimester developing human brain. bioRxiv 2022.10.24.513487 (2022) doi:10.1101/2022.10.24.513487.

30. Soldatov, R. et al. Spatiotemporal structure of cell fate decisions in murine neural crest. Science 364, (2019).

31. Petersen, C. P. & Reddien, P. W. Wnt signaling and the polarity of the primary body axis. Cell 139, 1056–1068 (2009).

32. Mulligan, K. A. & Cheyette, B. N. R. Wnt signaling in vertebrate neural development and function. J. Neuroimmune Pharmacol. 7, 774–787 (2012).

33. Liu, S. et al. Yap Promotes Noncanonical Wnt Signals From Cardiomyocytes for Heart Regeneration. Circ. Res. 129, 782–797 (2021).

34. Monroe, T. O. et al. YAP Partially Reprograms Chromatin Accessibility to Directly Induce Adult Cardiogenesis In Vivo. Dev. Cell 48, 765–779.e7 (2019).

35. Jiang, L., Li, J., Zhang, C., Shang, Y. & Lin, J. YAP-mediated crosstalk between the Wnt and Hippo signaling pathways (Review). Mol. Med. Rep. 22, 4101–4106 (2020).

36. Shalhout, S. Z. et al. YAP-dependent proliferation by a small molecule targeting annexin A2. Nat. Chem. Biol. 17, 767–775 (2021).

37. Laissue, P. P., Alghamdi, R. A., Tomancak, P., Reynaud, E. G. & Shroff, H. Assessing phototoxicity in live fluorescence imaging. Nat. Methods 14, 657–661 (2017).

38. Icha, J., Weber, M., Waters, J. C. & Norden, C. Phototoxicity in live fluorescence microscopy, and how to avoid it. Bioessays 39, (2017).

39. Hannezo, E. & Heisenberg, C.-P. Mechanochemical Feedback Loops in Development and Disease. Cell 178, 12–25 (2019).

40. Rifes, P. et al. Modeling neural tube development by differentiation of human embryonic stem cells in a microfluidic WNT gradient. Nat. Biotechnol. 38, 1265–1273 (2020).

41. Schindelin, J. et al. Fiji: an open-source platform for biological-image analysis. Nat. Methods 9, 676–682 (2012).

42. Pietzsch, T., Saalfeld, S., Preibisch, S. & Tomancak, P. BigDataViewer: visualization and processing for large image data sets. Nat. Methods 12, 481–483 (2015).

43. Kaya-Okur, H. S. et al. CUT&Tag for efficient epigenomic profiling of small samples and single cells. Nat. Commun. 10, 1930 (2019).

44. Zenk, F. et al. Analyzing the Genome-Wide Distribution of Histone Marks by CUT&Tag in Drosophila Embryos. Methods Mol. Biol. 2655, 1–17 (2023).

45. Langmead, B. & Salzberg, S. L. Fast gapped-read alignment with Bowtie 2. Nat. Methods 9, 357–359 (2012).

46. Ramírez, F. et al. deepTools2: a next generation web server for deepsequencing data analysis. Nucleic Acids Res. 44, W160–5 (2016).

47. Wahle, P. et al. Multimodal spatiotemporal phenotyping of human retinal organoid development. Nat. Biotechnol. 1–11 (2023).

48. Stuart, T. et al. Comprehensive Integration of Single-Cell Data. Cell 177, 1888–1902.e21 (2019).

49. He, Z., Brazovskaja, A., Ebert, S., Camp, J. G. & Treutlein, B. CSS: cluster similarity spectrum integration of single-cell genomics data. Genome Biol. 21, 224 (2020).

50. Krull, A., Buchholz, T.-O. & Jug, F. Noise2Void learning denoising from single noisy images. arXiv [cs.CV] 2129–2137 (2018).

51. Royer, L. A., Bragantini, J., Solak, A. C., Haase, R. & Yang, B. royerlab/dexp: First zenodo release. (2022) doi:10.5281/zenodo.7039059.

52. Virtanen, P. et al. SciPy 1.0: fundamental algorithms for scientific computing in Python. Nat. Methods 17, 261–272 (2020).

53. de Medeiros, G. et al. Multiscale light sheet organoid imaging framework. Nat. Commun. 13, 1–14 (2022).

54. Klein, S., Staring, M., Murphy, K., Viergever, M. A. & Pluim, J. P. W. elastix: a toolbox for intensity-based medical image registration. IEEE Trans. Med. Imaging 29, 196–205 (2010).

55. Shamonin, D. P. et al. Fast parallel image registration on CPU and GPU for diagnostic classification of Alzheimer’s disease. Front. Neuroinform. 7, 50 (2013).

56. Arganda-Carreras, I. et al. Trainable Weka Segmentation: a machine learning tool for microscopy pixel classification. Bioinformatics 33, 2424–2426 (2017).

57. Garreta, R. & Moncecchi, G. Learning scikit-learn: Machine Learning in Python. (Packt Publishing Ltd, 2013).

58. van der Walt, S. et al. scikit-image: image processing in Python. PeerJ 2, e453 (2014).

59. Attene, M. A lightweight approach to repairing digitized polygon meshes. Vis. Comput. 26, 1393–1406 (2010).

60. Dawson-Haggerty, M. et al. trimesh. https://trimsh.org/ (2019).

61. Arzt, M. et al. LABKIT: Labeling and Segmentation Toolkit for Big Image Data. Frontiers in Computer Science 4, (2022).

62. Yamauchi, K., Haase, R. & Sobolewski, P. morphometrics/morphometrics: v0.0.9. (2023) doi:10.5281/zenodo.8256767.

63. Wolf, F. A., Angerer, P. & Theis, F. J. SCANPY: large-scale single-cell gene expression data analysis. Genome Biol. 19, 15 (2018).

