## Supplementary material guide for "Morphodynamics of human early brain organoid development"

- **Supplementary Videos:** Ten supplementary videos are provided with the manuscript. Links to the videos as well as their legends are provided in the Supplementary videos pdf.
- **Supplementary methods table 1:** Supplementary methods table outlining details of the scRNAseq and lightsheet experiments done in this study.
- **Supplementary Table 1:** Gene list of pseudotime dependent genes from day 5 to day 11.
- **Supplementary Table 2:** Gene ontology term analysis list of pseudotime dependent genes.
- **Supplementary Table 3:** Differentially expressed genes between matrigel and no matrix organoids from an scRNAseq dataset sequenced on day 13.
- **Supplementary Table 4:** Gene ontology term analysis list for genes upregulated in matrigel (day 13).
- **Supplementary Table 5:** Gene ontology term analysis list for genes upregulated in no matrix (day 13).
- **Supplementary Table 6:** Differentially expressed genes between matrigel and no matrix organoids from an scRNAseq dataset sequenced on day 16.
- **Supplementary Table 7:** Differentially expressed genes between Control and YAP1 activator treated organoids from an scRNAseq dataset sequenced on day 10.
- **Supplementary Table 8:** Differentially expressed genes between Control and YAP1 activator treated organoids from an scRNAseq dataset sequenced on day 16.
