## Supplementary methods table 1 for "Morphodynamics of human early brain organoid development"

**Table 7:** Supplementary methods table

| Serial No. | Experiment dataset | Data type | Timepoint | Figure | Cell lines used | Time resolution |
|---|---|---|---|---|---|---|
| 1 | Organoid development timecourse (Protocol I) | scRNAseq | Days (5,7,11,16,21,30) | Figure 1,2 | Histone2B-mEGFP, | - |
| 2 | Matrix perturbation (Protocol I) | scRNAseq | Day 13 | Figure 4 | WTC-11 | - |
| 3 | YAP activator (Protocol I) | scRNAseq | Day 10 | Figure 4 | WTC-11 | - |
| 4 | Matrix perturbation (Protocol II) | scRNAseq | Day 16 | Extended Data Figure Figure 6 | WTC-11 | - |
| 5 | YAP activator (Protocol II) | scRNAseq | Day 16 | Extended Data Figure 6 | WTC-11 | - |
| 6 | Sparse and multi-mosaic organoids | Lightsheet | Days 4-12 | Figure 1,2,3 | WTC-11 90% + 2% each of: Histone2B-mEGFP , mEGFP-Beta-Actin, mTagRFP–T–CAAX, mTagRFP–T-TUBA1B, mTagRFP–T-LMNB1 | 30 minutes |
| 7 | Matrix perturbation (Protocol I) | Lightsheet | Days 4-12 | Figure 2,3 | WTC-11 94% + 2% each of: mEGFP-Beta-Actin, mTagRFP–T-TUBA1B, mTagRFP–T-LMNB1 | 60 minutes |
| 8 | YAP activator (Protocol I) | Lightsheet | Days 4-10 | Figure 4 | WTC-11 94% + 2% each of: mEGFP-Beta-Actin, mTagRFP–T-TUBA1B, mTagRFP–T-LMNB1 | 60 minutes |
| 9 | Mosaic organoid (Protocol I) | Lightsheet | Days 4-19 | Extended Data Figure 1 | WTC-11 (98%) + 1%: Histone2B-mEGFP, mTagRFP–T–CAAX, | 30 minutes |
| 10 | NKX2-1:GFP | Lightsheet | Days 0-21 | Extended Data Figure 1 | HES3 (NKX2-1:GFP) | 60 minutes |
