## Supplementary videos guide for "Morphodynamics of human early brain organoid development"

**Supplementary video 1: Sparse and multi-mosaic fluorescently labeled brain organoid imaged for one week.**

Link to video: https://polybox.ethz.ch/index.php/s/qQhJfvf4Ms2wSn0

Video shows maximum intensity projections of a developing sparse and multi-mosaic organoid brain from a time lapse dataset of 188 hour lightsheet imaging experiment. Organoids contain 5 different cell lines that contain stable genetic tagging of proteins with red or green fluorescent protein (RFP, GFP), as well as unlabeled cells, showing nuclear membrane (Lamin, RFP, magenta), plasma membrane label (CAAX, RFP, magenta), actin (GFP, green), tubulin (RFP, magenta), and nuclei (Histone, GFP, green). Timepoint 00:00:00 refers to the onset of imaging. Time is in hours and the scale bar is 100 micrometers.

**Supplementary video 2: 3D rotating visualization of a sparse and multi-mosaic organoid.**

Link to video: https://polybox.ethz.ch/index.php/s/XI1pBv9gyIh0y5D

3D z-stack rotations to visualize the dataset in supplementary video 1 at 93.5 hours (day 7) into imaging.

**Supplementary video 3: Lumen development, expansion and fusion events in early organoid development.**

Link to video: https://polybox.ethz.ch/index.php/s/PO6SU6ubR5QFwCx

Video shows cross-section of a developing organoid showing nuclear membrane (Lamin, RFP, orange), plasma membrane label (CAAX, RFP, orange), actin (GFP, cyan), tubulin (RFP, orange), and nuclei (Histone, GFP, cyan). The blue dot and line track shows fusion of two lumen. Time is in hours and the scale bar is 100 micrometers.

**Supplementary video 4: Organoid and lumen tissue dynamics in multiple brain organoids imaged in one experiment.**

Link to video: https://polybox.ethz.ch/index.php/s/wtqTLIuVO3Z3eM1

Video shows cross sections of 16 different organoids that were imaged together in one experiment. The segmented lumina are false colored with red hot LUT and the organoid tissue is shown in grayscale. Time is in hours and the scale bar is 500 micrometers.

**Supplementary video 5: 3D lumen morphodynamics in multiple brain organoids imaged in one experiment.**

Link to video: https://polybox.ethz.ch/index.php/s/nwoSMB2PdHIuJpq

Video shows 3D segmented lumen in 16 different organoids that were imaged together in one experiment. The experiment includes the organoids shown in supplementary video 1-4. The organoid outlines are shown as light colored contrast and the lumina are shown in cyan. Time is in hours and the scale bar is 500 micrometers.

**Supplementary video 6: Neuroectoderm development and lumen expansion dynamics in early organoid morphogenesis.**

Link to video: https://polybox.ethz.ch/index.php/s/f4dB0CsUIqT6VMq

Time lapse video showing cross-section of a developing lumen to show changes in lumen and neuroepithelium morphologies over time. The false coloring corresponds to nuclear membrane (Lamin, RFP, orange), plasma membrane label (CAAX, RFP, orange), actin (GFP, cyan), tubulin (RFP, orange), and nuclei (Histone, GFP, cyan). Time is in hours and the scale bar is 75 micrometers.

**Supplementary video 7: Developing organoids cultured in matrigel, without any extrinsic matrix and embedded in agarose.**

Link to video: https://polybox.ethz.ch/index.php/s/05wn9NhBehEXtYB

Time lapse video showing cross-sections of example neural organoids that were imaged in one experiment and were either embedded in matrigel, given no matrix or embedded in agarose. The false coloring corresponds to nuclear membrane (Lamin, RFP, orange), actin (GFP, cyan), tubulin (RFP, orange), Time is in hours and the scale bar is 100 micrometers.

**Supplementary video 8: Multi-sample parallel imaging of organoids grown in different extracellular matrix environments.**

Link to video: https://polybox.ethz.ch/index.php/s/b5vbBkBJ3qAnXgH

Time lapse video showing cross sections of 16 different organoids that were imaged together in one experiment and were either embedded in matrigel (first row), given no matrix (second and fourth row) or embedded in agarose (third row). The segmented lumina are false colored with red hot LUT and the organoid tissue is shown in grayscale. Time is in hours and the scale bar is 500 micrometers.

**Supplementary video 9: 3D lumen morphodynamics across multiple brain organoids cultured in matrigel, without any extrinsic matrix and embedded in agarose.**

Link to video: https://polybox.ethz.ch/index.php/s/8uvwYJUOzT1pNEp

Video shows 3D segmented lumen in 16 different organoids that were imaged together in one experiment. The experiment includes the organoids shown in supplementary video 7. The organoid outlines are shown as light colored contrast and the lumina are shown in cyan. The first row shows organoids embedded in matrigel, second and last row shows organoids imaged with no matrix and the third row shows organoids embedded in agarose. The false coloring corresponds to nuclear membrane (Lamin, RFP, orange), actin (GFP, cyan), tubulin (RFP, orange). Time is in hours and the scale bar is 500 micrometers.

**Supplementary video 10: Organoid developmental dynamics under control and YAP1 activator condition.**

Link to video: https://polybox.ethz.ch/index.php/s/TGLJIkGB8fmmj4K

Video shows maximum intensity projections of two different organoids from timepoint 00:72:00 hour (Day 8) onwards. Control is on the left and an organoid that was treated with YAP1 activator Py-60 on day 7 is on the right. The false coloring corresponds to nuclear membrane (Lamin, RFP, orange), actin (GFP, cyan) and tubulin (RFP, orange), Time is in hours and the scale bar is 100 micrometers.
